# The complete λ-carrageenan depolymerization cascade from a marine Pseudoalteromonad revealed by structural analysis of the enzymes

**DOI:** 10.1101/2025.01.09.632246

**Authors:** Chelsea J. Vickers, Andrew G. Hettle, Joanne K. Hobbs, Sarah Shapiro-Ward, Benjamin Pluvinage, Brendon J. Medley, Bailey E. McGuire, Liam Mihalynuk, Nitin, Wesley F. Zandberg, Alisdair B. Boraston

## Abstract

Carrageenans are a complex group of polysaccharides derived from the cell walls of red macroalgae. They are an abundant, yet recalcitrant nutrient source for most marine heterotrophic bacteria. Some member species of the *Pseudoalteromonas* genus are effective at metabolizing carrageenan. However, the enzymatic pathway for λ-carrageenan, one of the most sulfated naturally occurring polysaccharides, remains unknown. Using detailed structural analysis by X-ray crystallography we reveal the sophisticated and cyclic enzymatic cascade deployed by *Pseudoalteromonas distincta* (strain U2A) to utilize λ-carrageenan. The cascade incorporates ten glycoside hydrolases and five sulfatases that are specific for λ-carrageenan and cooperate to completely deconstruct this polysaccharide, thus yielding galactose monosaccharides for subsequent energy production. The detailed molecular understanding of λ-carrageenan depolymerisation provided includes structural evidence for a lesser described sulfatase catalytic mechanism and elucidation of a distinct catabolic cascade that is unique from previously described carrageenan metabolic pathways. This insight also holds potential for the application of enzymatic logic in the generation of high value products from abundant natural biopolymers, such as carrageenans.

## Introduction

Carrageenan is a family of linear sulfated polysaccharides found in some red seaweeds. They belong to a class of marine based anionic polymers/hydrocolloids, including agars, that are well known for their gelling properties and are used in a variety of laboratory and industrial applications (1–3). They also represent an abundant, yet recalcitrant nutrient source for marine heterotrophic bacteria with algae derived polysaccharides constituting up to 35% of bioavailable carbon in the ocean (4–6). The molecular structure of carrageenan is characterized by repeating units of disaccharides comprising D-galactose with these sugar units connected by alternating α-1,3 and β-1,4 glycosidic linkages. The specific type of carrageenan is defined by whether the second D-galactose of the disaccharide unit (i.e. the residue that is α-linked to the preceding residue) has a 3,6-anhdyro configuration and the specific sulfation pattern. The most common carrageenan types are κ-, ι-, and λ-carrageenan. κ- and ι-carrageenan both possess the 3,6-anhydro-D-galactose with 4-O-sulfo-D-galactose as the first residue in the repeating unit. In λ-carrageenan the 3,6-anhydro-D-galactose is replaced by D-galactose and also has a 2-O-sulfate modification. λ-carrageenan lacks 3,6-anhydro-D-galactose and instead has a β-1,4-linked repeating disaccharide of α-1,3- linked 2-O-sulfo-D-galactose (G2S) and 2,6-di-O-sulfo-D-galactose (G2,6S). The expected ratio of 1.5 sulfate groups per monosaccharide makes λ-carrageenan one of the most highly sulfated polysaccharides presently known to exist (7).

The sulfation of polysaccharides presents a challenge to the bacteria that would metabolize them because, at some point in the metabolic pathway, additional steps to remove the sulfate groups must be taken. This has been elegantly illustrated by, for example, the complete determination of pathways dedicated to glycosaminoglycan and porphyran metabolism by gut bacteria, and κ-/ι- carrageenan metabolism by marine bacteria (8–12). Given its extraordinary sulfation level, λ- carrageenan may present a unique challenge to microbes seeking to use it as a carbon source. Indeed, the observation of a growth phenotype on λ-carrageenan that is potentially linked to a genetic locus encoding the relevant enzymatic machinery is quite rare, and at present most common to the genus *Pseudoalteromonas* (10, 12–15). This is a genus of marine bacteria known for its diversity and prevalence in various marine environments, and for its diverse metabolic assimilation of organic matter, particularly algal polysaccharides (16–20).

With respect to λ-carrageenan metabolism, CglA (PCAR9_p0052) from *Pseudoalteromonas carrageenovora* 9^T^ and its ortholog in *Pseuodalteromonas* bacterium strain CL19 are λ- carrageenases. These are classified into glycoside hydrolase family 150 (GH150) and characterized as *endo*-hydrolases that hydrolyze the β-1,4-glycosidic linkages of λ-carrageenan in a non-processive fashion (20, 21). More recently, genome sequencing of *P. carrageenovora* 9^T^ identified a large genetic locus dedicated to carrageenan metabolism, which we refer to as a **<u>car</u>**rageenan specific **<u>p</u>**olysaccharide **<u>u</u>**tilization **l**ocus (CarPUL) (22). Notably, the gene encoding a CglA is present in the CarPUL. Other proteins encoded in the CarPUL were hypothesized to contribute to the growth phenotype of *P. carrageenovora* 9^T^ on λ-carrageenan, though the molecular mechanisms by which bacteria deconstruct this complex polysaccharide remain unknown.

In this study, we report the ability of *Pseudoalteromonas distincta* (strain U2A, herein referred to only as U2A) isolated from red macroalgae found in the intertidal zone of south Vancouver Island, Canada (10), to grow on λ-carrageenan. Through integrated analysis of ten enzymes encoded in a polysaccharide utilization locus we refer to as a λ-CarPUL, which is syntenic to a region of the *P. carrageenovora* 9^T^ CarPUL, we link this specific locus to the ability of U2A to metabolize this highly sulfated polysaccharide. Using a purified λ-neocarratetraose oligosaccharide we determined the structures of the final eight of the enzymes in the pathway trapped in complex with their respective substrates (or in one case an inhibitor analogue), which were rationally generated by sequential treatment of the oligosaccharide by the enzymes upstream in the cascade. This data, supported by enzyme assays, provides unprecedented insight into the molecular details of substrate recognition and turnover for a complete and sophisticated enzyme cascade, and is the first such insight for system performing λ-carrageenan depolymerization. This provides an opportunity to harness this enzymatic logic to enhance biosynthetic platforms for generation of high value products from abundant marine resources such as red algae and other producers of industry relevant hydrocolloids.

## Results

### PUL structure and bacterial growth

We previously reported the isolation and genome sequencing of five *Pseudoaltermonas* strains isolated from the surface of *Fucus* sp. obtained from the same marine environment(10). *Pseudoalteromonas fuliginea* PS47 (referred to herein as PS47) and *P. distincta* U2A both harbored plasmids encoding a CarPUL, and both displayed the ability to grow on κ- and ι-carrageenan. The U2A CarPUL, however, includes an additional segment that is syntenic to a region of the *Pseudoalteromonas carrageenovora* 9^T^ CarPUL (Fig. 1A). This additional segment of the locus, which we refer to as the λ-CarPUL, was previously hypothesized by Gobet *et. al.* (2018) to confer λ-carrageenan metabolism to *P. carrageenovora* 9^T^ (22). Consistent with their lack of a λ-CarPUL, PS47 and *P. fuliginea* PS42 (referred to as PS42), the latter of which completely lacks a CarPUL, failed to display any growth in liquid culture containing λ-carrageenan as a sole carbon source (Fig. 1B). The PS47 isolate possesses the ι/κ-CarPUL, as previously reported (10), yet this does not seem to confer any ability to utilize λ-carrageenan. However, growth was observed for U2A in conditions with 0.5% λ-carrageenan as the sole carbon source (Fig. 1B). All strains grew on 0.4% galactose, which was used as a positive control. To assess metabolic localisation U2A was grown in liquid medium with 0.2 % λ-carrageenan and then separated into extracellular (ECF, or cleared culture supernatant) and total cell fractions (TCF, lysed cells removed from culture medium). These fractions were then incubated with fresh λ-carrageenan at 1%. Consistent with the ability of U2A to utilize λ-carrageenan, both cellular fractions showed depolymerization of λ-carrageenan, though with different product profiles (Fig. 1C and 1D). The TCF produced a product with a mobility the same as that of the galactose standard, providing evidence of the enzyme machinery necessary to depolymerize λ-carrageenan to galactose. To provide a link between the presence of genes encoding a putative λ-carrageenan pathway and the ability of U2A to grow on this substrate we pursued biochemical characterization of enzymes encoded by the λ-CarPUL.

**Figure 1.**
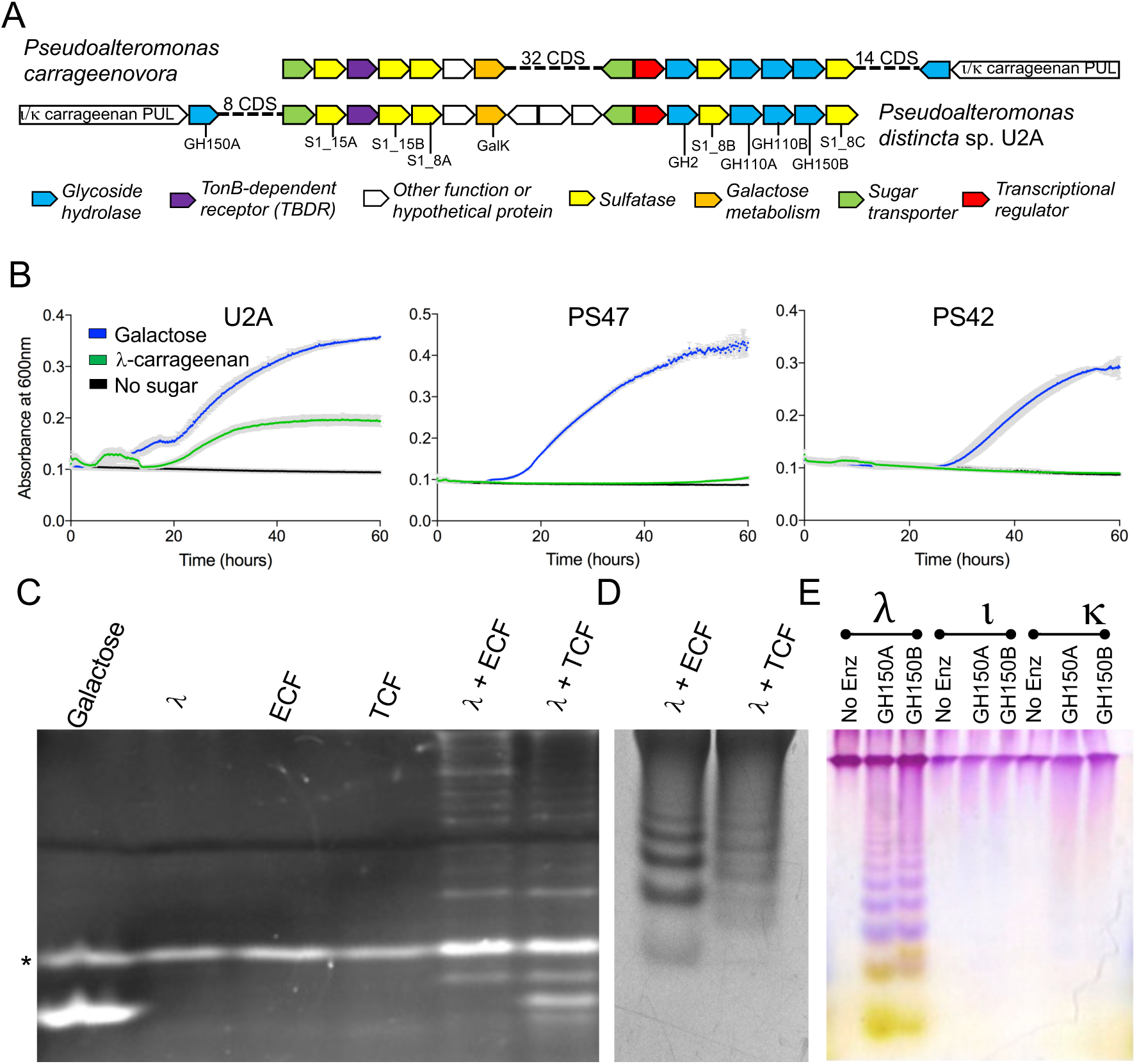
*Pseudoalteromonas distincta* U2A λ-CarPUL organization and λ-carrageenan growth experiments. A) Schematic of the additional λ-CarPUL segment in the *Pseudoalteromonas distincta* U2A (U2A) CarPUL that is syntenic to a region of the *Pseudoalteromonas carrageenovora* 9^T^ CarPUL and is hypothesised to confer λ-carrageenan metabolism. Each gene is represented by an arrow, which are colour coded according to putative function. B) Carrageenan growth experiments of isolated *Pseudoalteromonas* sp where growth on λ-carrageenan is exhibited by U2A exclusively. Gray bars indicate the error from *n* = 4 independent experiments. FACE (C) and C-PAGE (D) metabolic localization experiments showing different λ-carrageenan depolymerisation profiles by U2A extracellular fractions (ECF or cleared culture supernatant) and total cell fractions (TCF, lysed cells removed from culture medium). E) U2A C-PAGE carrageenan activity experiment showing different depolymerisation profiles of GH150A and GH105B exclusively on λ-carrageenan. The asterisk in panel B indicates the free label ANTS.

### Initial λ-carrageenan depolymerization

The λ-CarPUL of U2A encodes two putative family 150 GHs, which we refer to as GH150A (locus tag EU511_08665) and GH150B (EU511_08535). GH150A and GH150B have amino acid sequence identities of 99% and 26%, respectively, with CglA (PCAR9_p0052) from *P. carrageenovora* 9^T^ and the CglA ortholog from *Pseuodalteromonas* bacterium strain CL19 (20, 21). Both previously characterized λ-carrageenases are classified as λ-carrageenan-specific *endo*-hydrolases that hydrolyze the β-1,4-glycosidic linkage in a non-processive fashion. Through fusion of a maltose binding protein (MBP) at the N-terminus of the GH150 proteins we were able to produce and purify small amounts of the MBP fusions as soluble proteins. Activity of the MBP-GH150 fusions on λ-carrageenan produced ladders of products but neither enzyme was active on κ- or ι-carrageenan (Fig. 1E). The products produced by GH150B displayed a small but reproducible upward shift in mobility relative to those of GH150A, indicating a slightly different set of products. Given that GH150A is essentially identical to CglA from *P. carrageenovora* 9^T^, which has been shown to produce a fully sulfated series of even numbered λ- neocarrageenoligosaccharides, it is reasonable to assume that GH150A produces the same series of products. To provide some insight into the potential differences in the GH150B products we purified the λ-neocarratetraose (λ−NC4) fraction and characterized it by CE-LIF and LC-MS (see Fig. S1 and Table S1 for characterization of the oligosaccharide). Multiple species were detected by both approaches with the majority of the sample (>80%) interpreted from the CE-LIF to be tetrasaccharide. LC-MS showed the tetrasaccharide to be in three differently sulfated forms of tetrasaccharide with six, five, and four sulfate groups (Hex_4_S_6_, Hex_4_S_5_, and Hex_4_S_4_, respectively), with the fully sulfated Hex_4_S_6_ predominating (∼70% of the tetrasaccharide population based on summed MS peak areas). The ability of GH150B to release incompletely sulfated λ- neocarrageenoligosaccharides distinguishes it from CglA, and likely explains the different product pattern of GH150B cf. GH150 revealed by C-PAGE (Fig. 1E) while suggesting that GH150B may be capable of cleaving in regions of λ-carrageenan that are not fully sulfated. We attempted to crystallize GH150A and GH150B but were unsuccessful. Therefore, we compared models generated with AlphaFold2 (23), which indicated an overall fused 3-domain architecture for both proteins (Fig. 2A and Fig. S2). This analysis revealed a patch of surface residues that is conserved amongst GH150 family members. This presumably comprises the catalytic machinery, or other functionally relevant residues, and resides in a deep cleft running the width of the protein that is created by the two lobes making up the protein (Fig. 2B). This region spans ∼12 Å of the cleft, or the approximate length of a disaccharide unit, possibly suggesting, by the level of conservation and dimensions, the location of the +1 and -1 catalytic subsites.

**Figure 2.**
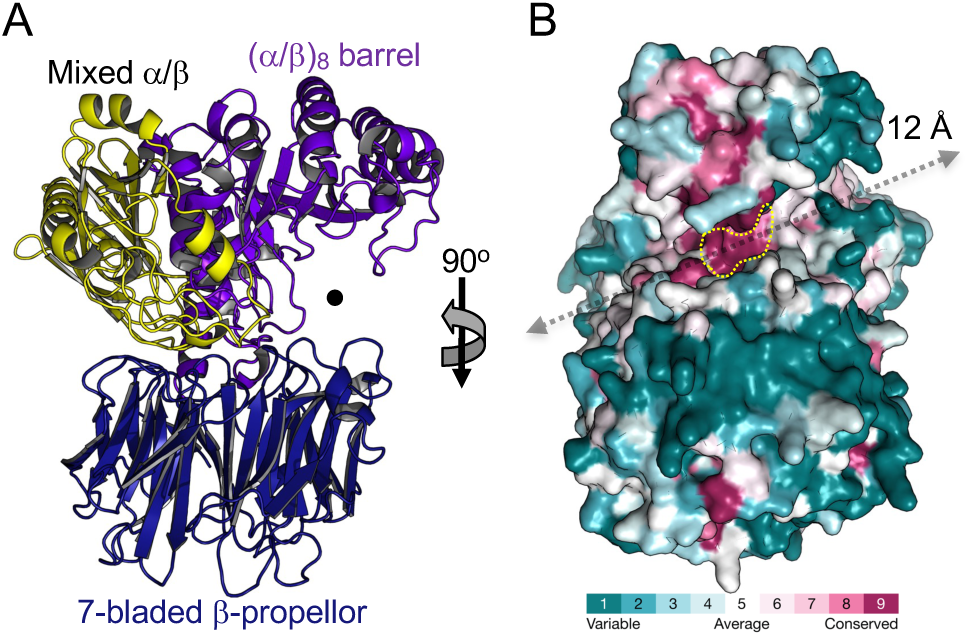
Structural modelling of GH150A and GH150B with AlphaFold 2. A) Cartoon representation of the GH150A AlphaFold 2 model as representative of the overall fold of both GH150A and GH150B (and other GH150 enzymes). The structure is coloured and labelled according to the three identified domains. B) Consurf analysis revealing conserved deep cleft running the width of the protein that is created by the two lobes. The length (approximately 12 Å) and location of the cleft is indicated by a grey arrow. The most conserved patch, which presumably comprised the catalytic machinery or other functionally relevant residues, is highlighted by the yellow dotted line.

### Identification of a λ-neocarrageenoligosaccharide specific exo-galactose-2[,6]-sulfate-2-O- sulfohydrolase

We previously characterized GH110B (EU511_08540) from the λ-CarPUL and identified it to be an *exo*-α-D-galactosidase that recognizes a non-sulfated galactose residue at the non-reducing terminus of α-1,3-galactobiose (αG2)(24). Given this activity we hypothesized that the 2S and/or 6S groups of the non-reducing terminal α-linked galactose in λ- neocarrageenoligosaccharides produced by the GH150 enzymes would have to be removed prior to GH110 activity. We used C-PAGE to separately screen the potential activity of all five λ-CarPUL encoded sulfatases, which were produced under conditions to promote their maturation to an active form, on the pool of oligosaccharides produced by the depolymerization of λ-carrageenan by GH150B (Fig. 3A). This revealed a slight but consistent upward shift in the mobility of the resulting oligosaccharides only in the presence of S1_8B, indicating an alteration to the oligosaccharide population by this putative sulfatase. To probe this in more detail we pursued structural analysis of S1_8B in complex with our preparation of λ−NC4 to provide additional insight into its activity.

**Figure 3.**
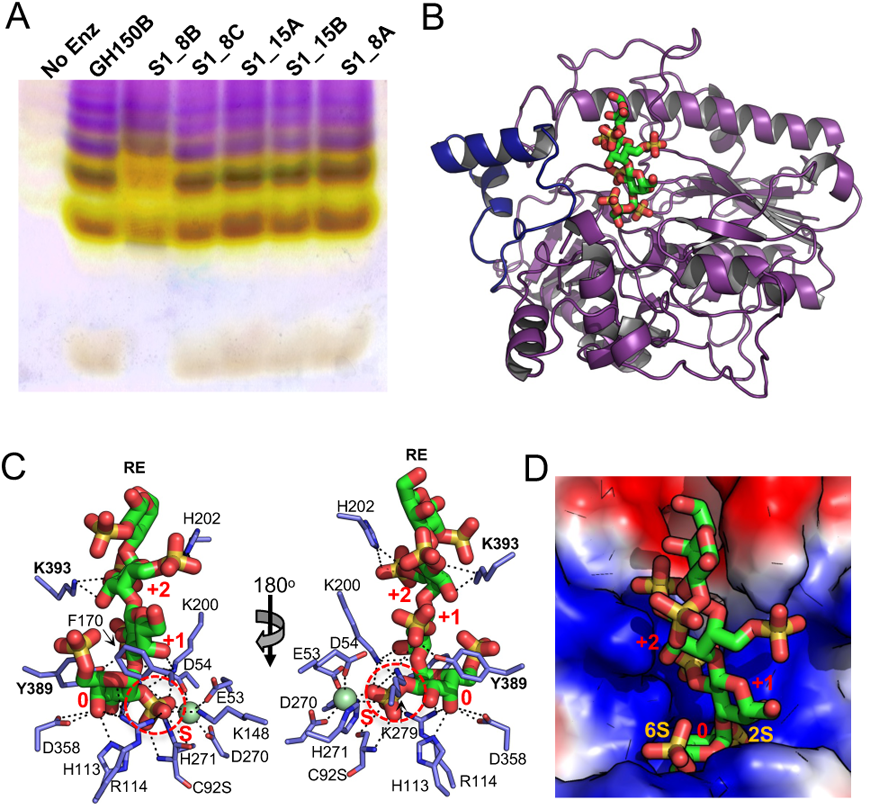
Characterisation of S1_8B. A) Screening of the five sulfatases in the U2A λ-CarPUL by C-PAGE to identify the λ-neocarrageenoligosaccharide specific exo-galactose-2[,6]-sulfate-2-O-sulfohydrolase. The substrates used for this screen was a pool of oligosaccharides produced by the depolymerization of λ-carrageenan by GH150B. S1_8B was the only sulfatase that showed activity on these oligosaccharides. B) The apo X-ray crystallographic structure of S1_8B (purple) revealing the typical two-domain alkaline phosphatase fold of the S1 family. The large active-site adjacent disordered loop that could only be modelled into the inactive C92S structure bound to λ-NC4 is shown in blue. C) Interactions between residues on the dynamic active site-adjacent loop and λ-NC4. The 2S of the non-reducing end G2,6S engages with the catalytic machinery in the S-subsite. D) Surface electrostatics show the catalytic machinery in the S-subsite is buried in the base of a positively charged active site pocket that is complementary to the negatively charged l-NC4 substrate.

The uncomplexed structure of S1_8B revealed the typical two-domain alkaline phosphatase fold of the S1 family (Fig. 2B). However, a relatively large loop comprising approximately 40 residues (∼ residues 380 to 420) adjacent to the active site was disordered and could not be modelled. Subsequently, an inactive C92S mutant was used to generate crystals that were soaked with the λ−NC4 oligosaccharide. The complexed structure determined to 2.1 Å resolution was solved with a tetrasaccharide occupying the active site and the disordered loop regions in each molecule in the asymmetric unit were able to be built in (Fig. 3B). We were able to model both tetrasaccharides as fully sulfated λ−NC4 (*i.e.* Hex_4_S_6_)(Fig. S3A) and show that two residues (Y389 and K393) from the dynamic loop directly interact with the bound tetrasaccharide at subsite +1 and +2, respectively (Fig. 3C). The mode of oligosaccharide binding had the 2S of the non-reducing end G2,6S engaging the catalytic machinery in the S-subsite (Fig. 3C), which was buried in the base of the complementary positively charged active site pocket (Fig. 3D). The 6S group did not make any additional specific interactions in the 0-subsite, suggesting this sulfate group may be dispensable for substrate recognition. Though we were able to model all four monosaccharides of the tetrasaccharide, which would potentially indicate the presence of up to a +3 subsite, only specific interactions in +1 and +2 subsites were evident. This included hydrogen bonds with the 2S groups of both monosaccharides in these subsites (Fig. 3C).

Overall, this structural analysis of S1_8B indicates *exo*-galactose-2,6-sulfate-2-O-sulfohydrolase activity for this enzyme; however, it is possible that the 6S is not strictly required for substrate recognition. The structural details of substrate recognition for other members of the S1_8 subfamily is presently unknown, but the structural assignment of this 2-O-sulfohydrolase specificity to S1_8B is consistent with the *exo*-xylose-2-sulfate-2-O-sulfohydrolase and N-sulfoglucosamine sulfohydrolase activities of other S1_8 family members (25, 26).

### Hydrolysis of the terminal α-linked galactose is dependent upon 2-O-desulfation

GH110A (EU511_08545) displayed no activity on *p*NP-α-D-galactopyranoside (*p*NP-α-Gal) or αG2 (Fig. S4A), making it unlike GH110B, which was active on both substrates, and raising the question of whether it may accommodate a sulfated substrate (24). We assessed the activity of GH150B and GH110A together on λ-carrageenan, as well as both of these enzymes in combination with each of the individual sulfatases. Only when S1_8B was included was there a change in the products (Fig. 4A). Furthermore, the product profile was different than when only GH150B and S1_8B was used to digest λ-carrageenan (Fig. 3A *cf.* Fig. 4A). Consistent with this, we used the galactose release assay, which also detects G6S (Fig. S4B), to show that only the GH150B, GH110A, and S1_8B combination resulted in detectable monosaccharide release from the pool of GH150B generated λ-neocarrageenoligosaccharides (Fig. 4B). These results indicate that removal of the 2-sulfate from the non-reducing terminal galactose by S1_8B can initiate the depolymerization of λ-neocarrageenoligosaccharides by GH110A.

**Figure 4.**
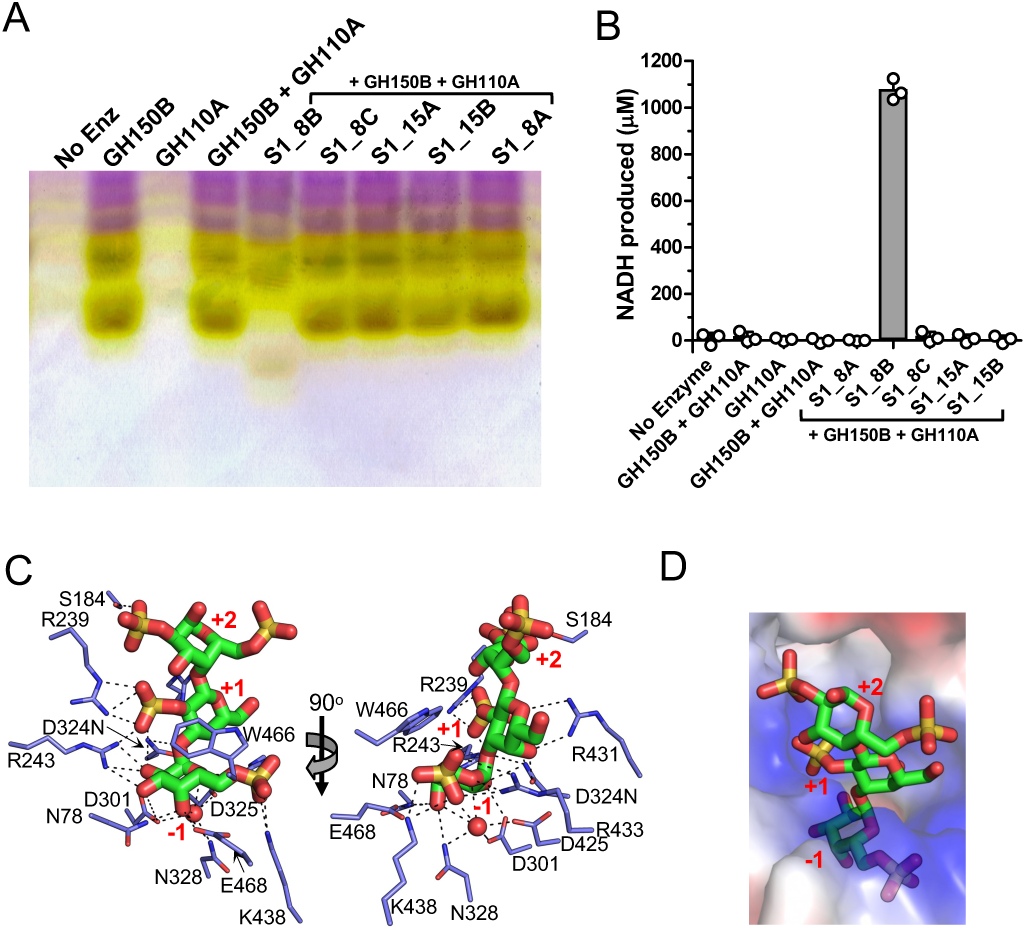
Characterisation of GH110A. A) C-PAGE screening for GH110A and sulfatase activities in combination on GH150B produced λ-carrageenan oligosaccharides. B) Galactose release assays confirm monosaccharide release from assays containing GH150B, S1_8B and GH110A indicating removal of G6S from the λ-neocarrageenoligosaccharides. Grey bars indicate the mean values of triplicate independent measurements and the error bars the standard deviation. Closed circles show individual data points. C) X-ray crystallographic structure of GH110A D324N mutant showing the active site bound to λ-NC4 that was pre-treated with S1_8B. D) Surface electrostatics show complementary charge of the +1 and +2 subsites to the edge of the glycan region that presents the sugars with 2S modifications.

Based on prior knowledge of the catalytic residues in GH110B and the structure of GH110A with G6S (Figs. S3B, S5A, and S5B), we generated an inactive GH110A D324N mutant and explored the molecular basis of substrate recognition by determining its structure in complex with λ−NC4 that was pre-treated with S1_8B. Though the F_o_-F_c_ electron density maps suggested partial occupancy of sugar residues in the +1 and +2 subsites, possibly due to partial hydrolysis of the substrate, we were still able to model a trisaccharide into the active sites of each monomer in the asymmetric unit (Fig. 4C and Fig. S3C). The G6S fit tightly into -1 subsite in the same manner as observed in the G6S complex, including recognition of the 6S group (Fig. 4C and D). +1 and +2 subsites provide additional interactions with the oligosaccharide, though mainly in the +1 subsite. The electrostatic charge of the active site is notably complementary to the edge of the glycan presenting the 2S modifications on the sugars in the +1 and +2 subsites (Fig. 4D). These structure-activity results indicate that GH110A is an *exo*-6-sulfo-α-D-galactopyranosidase with at least three active site subsites.

### Identification of a λ-carrageenoligosaccharide active exo-β-D-galactopyranosidase

The activities of S1_8B and the GH110A generate a λ-carrageenoligosaccharide with a non-reducing end β- linked G2S residue. Given the frequent association of GH2 enzymes with β-galactosidase activity and the 2-O-sulfohydrolase activity of S1_8 enzymes we focused investigation of λ- carrageenoligosaccharide depolymerization by the GH2, in combination with the additional S1_8A, and S1_8C sulfatases in the CarPUL. GH2 was active on *p*NP-β-D-galactopyranoside (*p*NP-β- Gal; not shown) and Galβ1–4Gal (βG2; Fig. 5A), confirming the β-galactosidase activity of the protein. Using a galactose release assay, GH2 was not able to release additional detectable monosaccharide from the pool of GH150 generated λ-neocarrageenoligosaccharides that were first pre-treated with GH110A and S1_8B to generate λ-carrageenoligosaccharides. When S1_8C, but not S1_8A, was included in the assay a large increase in released monosaccharide was observed (Fig. 5B). This suggested that S1_8C removed the 2S modification from a non-reducing end β- linked G2S residue to produce an unsulfated non-reducing end β-linked galactose, thus allowing activity of the GH2 β-galactosidase.

**Figure 5.**
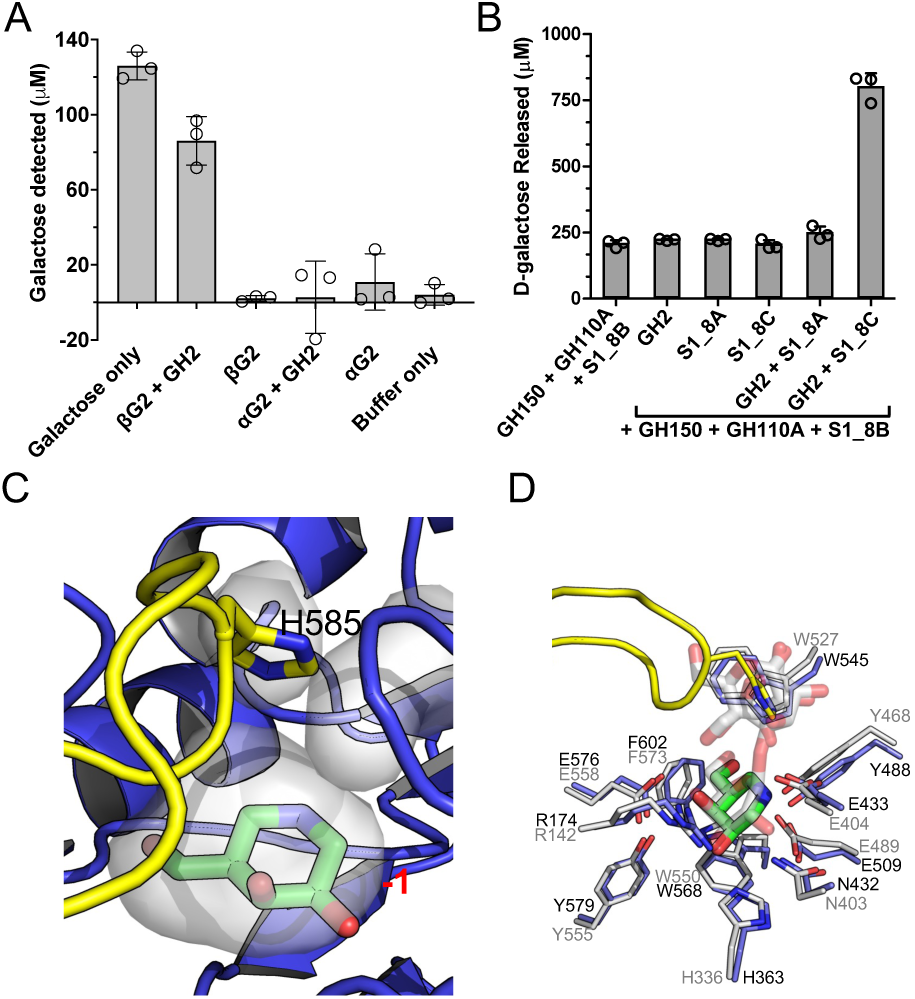
Release of unsulfated galactose by S1_8C and GH2. A) GH2 is a β-galactosidase that shows activity on Galβ1–4Gal in a galactose release assay. B) In galactose release assays GH2 releases monosaccharide from oligosaccharides produced from λ-carrageenan treated with a combination of GH150B, S1_8B and GH110A but only when S1_8C is included in assays i.e., S1_8C and GH2 work together to remove a monosaccharides from carrageenoligosaccharides with a terminal G2S at the non-reducing end. In panels A) and B) grey bars indicate the mean values of triplicate independent measurements and the error bars the standard deviation. Closed circles show individual data points. C) GH2 crystallographic structure showing the -1 subsite bound to inhibitor galactoisofagomine (GIF) with capping loop residue H585 at its apex immediately over the actives site. The -1 subsite of GH2 is contoured very closely around the GIF molecule such that a sulfated galactose residue could not be accommodated. D) Superimposition of the GH2 structure and β-galactosidase II from *Bacillus circulans* (PDB ID 7CWD).

The structure of GH2 in complex with the inhibitor galactoisofagomine (GIF) revealed the typical GH2 (β/α)_8_ barrel fold (Figs. S3C and S5C) with a pocket-like active site, which is capped by a loop with H585 at its apex immediately over the active site (Fig. 5C). The conformation of this loop effectively closes off the active site. This suggests that the inhibited form that was crystallized possesses a loop conformation that is not permissive to activity and that the loop must be mobile to allow substrate access. Consistent with this hypothesis, the B-factors of the capping loop are elevated relative to surrounding regions, indicating a degree of mobility (Fig. S5B). The -1 subsite is contoured very closely around the GIF molecule such that a sulfated galactose residue could not be accommodated (Fig. 5C). Indeed, the -1 subsite of GH2 is completely conserved with the active site of β-galactosidase II from *Bacillus circulans*, which is active on unmodified β-1,4- galactooligosaccharides (Fig. 5D) further indicating that GH2 is an *exo*-β-D-galactopyranosidase.

### Identification of a λ-carrageenoligosaccharide active exo-galactose-2-sulfate-2-O-sulfohydrolase

The dependency of GH2 activity on S1_8C to enable the removal of the terminal β-linked galactose from λ-carrageenoligosaccharides is most consistent with S1_8C as having λ- carrageenoligosaccharide *exo*-galactose-2-sulfate-2-O-sulfohydrolase activity. To provide additional support for this we pursued structural analysis of S1_8C in complex with λ-C3 (λ−carratriose or G2S-G2,6S-G2S), which was produced by pre-treatment of λ−NC4 with GH110A and S1_8B.

The unliganded structure of S1_8C comprising two monomers in the asymmetric unit was solved by molecular replacement using the model of S1_8B, with which it shares 30% amino acid sequence identity and a structural RMSD of 1.9 Å (Fig. 6A). The main structural differences occur in the C-terminal domain, which does not appear to be involved in the formation of the active site in either enzyme. A C68S mutation was introduced into S1_8C to render it inactive and crystals of this protein were soaked with λ-C3. We were able to model a disaccharide comprising λ−C2 in the active site of one monomer and G2S in the active site of the other monomer (Fig. S3D). λ−C2 makes an extensive series of interactions with the enzyme, specifically with the 2S group of the G2S residue engaging the catalytic machinery (Fig. 6B) in a blind-pocket active site (Fig. 6C). This supports assignment of S1_8C as another λ-carrageenoligosaccharide active *exo*-galactose-2- sulfate-2-O-sulfohydrolase in this CarPUL. The S and 0 subsites of S1_8C are very well conserved with those of S1_8B (Fig. 6D). The recognition of a terminal β-linked residue by S1_8C vs a terminal α-linked G2,6S by S1_8B results in different trajectory of substrate from the active site, which is enabled by very different plus (+) subsites in the two enzymes (Fig. 6E).

**Figure 6.**
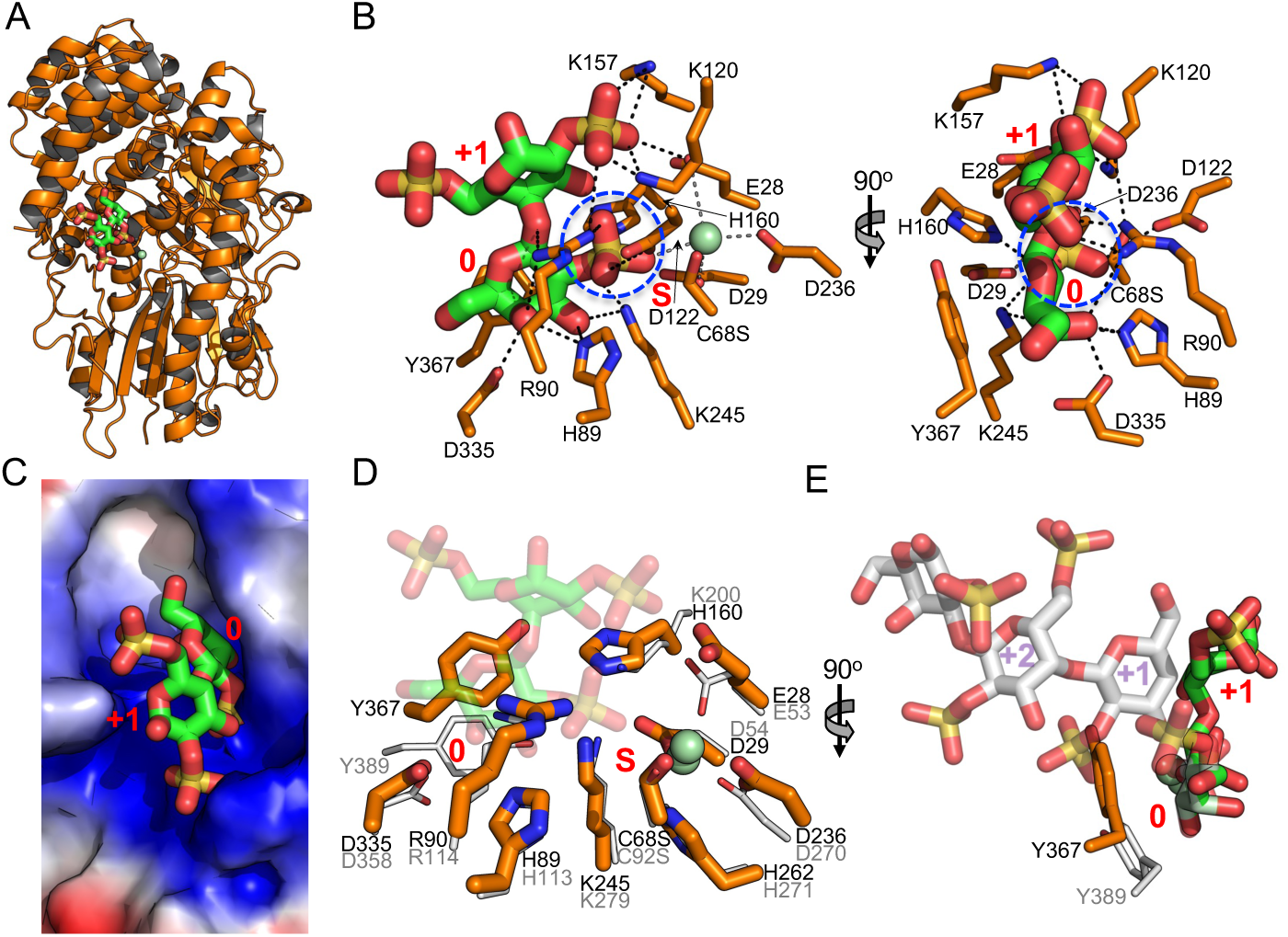
Characterisation of S1_8C. A) X-ray crystallographic structure of S1_8C. B) The S1_8C C68S structure bound to λ-C3 which makes extensive interactions with the active site, specifically with the 2S group of the G2S residue engaging the catalytic machinery. C) Solvent accessible surface showing a blind-pocket active site architecture. Superimposition of S1_8C and S1_8B showing D) conservation of the S and 0 subsites and E) differences in +1 and +2 subsites associated with the different trajectories of substrates when bound in active sites.

### Identification of exo-galactose-6-sulfate-6-O-sulfohydrolases

Given that the two characterized S1_8 sulfatases had 2S specificity, we reasoned that the S1_15 enzymes would be 6-sulfate-6-O-sulfohydrolases. Supporting this supposition, other S1_15 family members, in particular BT3333 from *Bacteroides thetaiotaomicron*, have N-acetylgalactosamine-6-sulfate exo-6-O-sulfohydrolase activity (i.e. 6S sulfatase activity)(27). We determined a preliminary apo structure of S1_15A by SAD phasing on an iodide derivative and then used the preliminary coordinates to solve the structure of an inactive S1_15A C94S mutant in complex with G6S (Fig. S3F). Subsequently we solved the structure of an S1_15B C84S mutant, also in complex with G6S (Fig. S3G). The structures were remarkably similar with significant variation only evident in a region near the entrance to the active site, identified by tryptophan residues W471 in S1_15A and W462 in S1_15B (Fig. 7A). For both enzymes, the interactions of G6S within the active sites were essentially identical and consistent with the specific interactions of galactose-6-sulfate-6-O-sulfohydrolases (Fig. 7B). In S1_15A the G6S residue was nearly completely enclosed in the active site by virtue of the α-helix containing W471 closing in over the active site (Fig. 7C and 7D). W471 then provides a platform against which the pyranose ring of the G6S packs. The enclosed active site would preclude recognition of an oligosaccharide. In contrast, the analogous helix in S1_15B is retracted from the entrance to the active site (Fig. 7A), providing an opening into the pocket (Fig. 7E and 7F). In neither case was there room to accommodate a 2S modification on the galactose in the 0-subsite.

**Figure 7.**
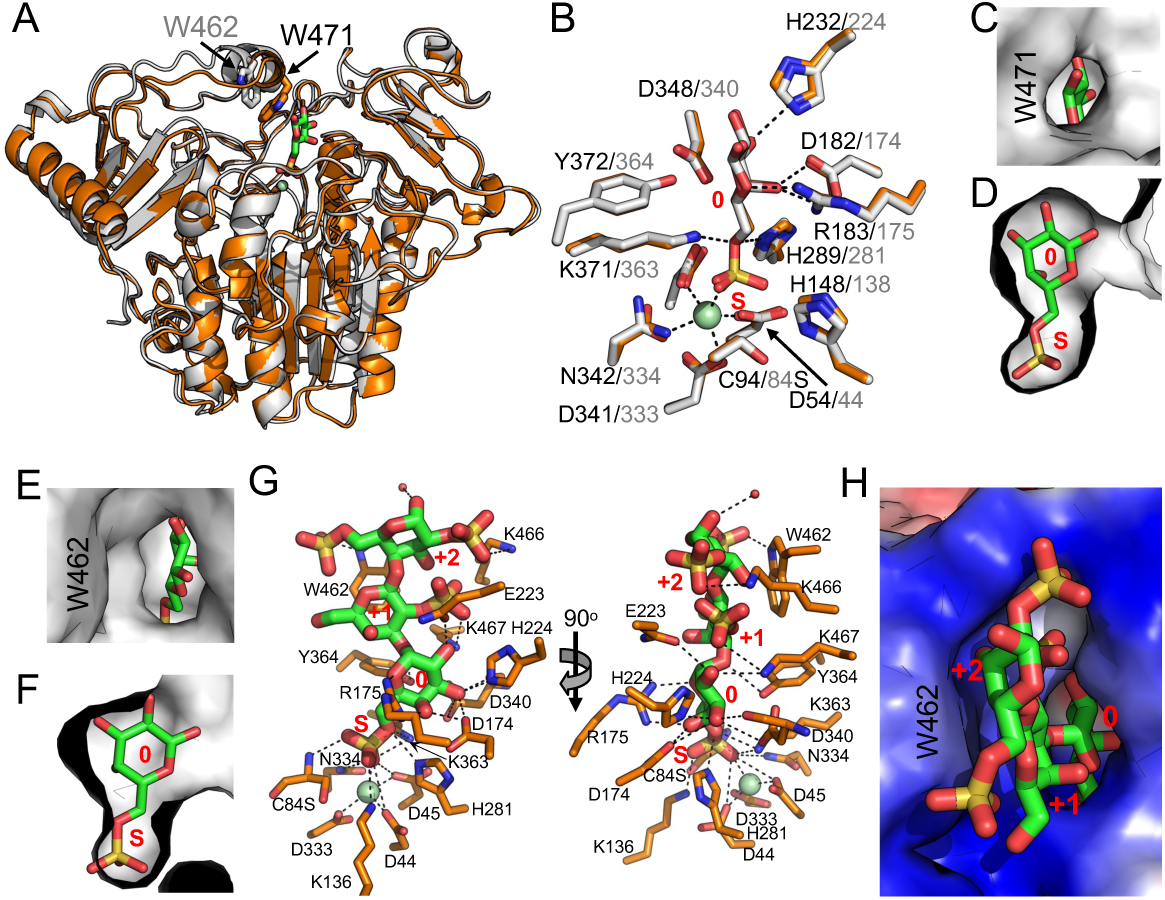
Structural analysis of 6S sulfatases S1_15A and S1_15B. A) Superimposition of S1_15A (orange) and S1_15B (grey) X-ray crystallographic structures showing structural variation at the entrance to the active site at tryptophans W471 and W462. B) Both enzymes interact with G6S in an almost identical manner consistent with the specific interactions of galactose-6-sulfate-6-O-sulfohydrolases. C and D) In S1_15A the G6S residue is nearly completely enclosed in the active site by virtue of the α-helix containing W471 closing in over the active site E and F) In contrast, the analogous helix in S1_15B is retracted from the entrance to the active site providing an opening into the pocket. In neither case was there room to accommodate a 2S modification on the galactose in the 0-subsite. G) S1_15B C84S in complex with λ-NC4 pretreated with S1_8B showing the poise of the terminal G6S in the active site was the same as that observed in the monosaccharide G6S complex. H) W462, which lines the entrance to the active site, packs against the pyranose ring of the G2S residue in the +1 subsite guiding the oligosaccharide into the base of the active site pocket.

These observations led us to propose that S1_15B would be an oligosaccharide active *exo-* galactose-6-sulfate-6-O-sulfohydrolase. To interrogate this, we solved the structure of S1_15B C84S in complex with λ−NC4 that had been pretreated with S1_8B to convert the non-reducing end residue to G6S. We were able to model a trisaccharide comprising G6S-G2S-G2,6S into the active site (Fig. S3H) . The poise of the terminal G6S in the active site was the same as that observed in the monosaccharide G6S complex (Fig. 6G). W462, which lines the entrance to the active site, packed against the pyranose ring of the G2S residue in the +1 subsite guiding the oligosaccharide into the base of the active site pocket (Fig. 6H). On the basis of these structural results, we assigned S1_15A as a monosaccharide (i.e. G6S) specific *exo-*galactose-6-sulfate-6- O-sulfohydrolase and S1_15B as a λ-neocarrageenoligosaccharide specific *exo-*galactose-6- sulfate-6-O-sulfohydrolase that obligately works after S1_8B.

### GH110B is a λ-neocarrageenoligosaccharide specific α-galactosidase

Our prior results indicated that the -1 subsite of GH110B has a complementary fit to an unsulfated sugar and has activity as an α-1,3-galactosidase, indicating that the activity of this GH would occur after S1_8B and S1_15B activity(24). Using a galactose release assay, we found that GH110B required at least S1_8B pre- treatment and that, under the conditions used, S1_15B did not provide the expected potentiation of activity (Fig. 8A). To provide insight into this we attempted to produce structures of an inactive GH110B D344N mutant in complex with substrates having a 6S on its non-reducing terminal galactose generated by S1_8B activity, as we did with GH110A, but we could not. However, we did obtain a complex with λ-NC4 that was pretreated with S1_8B and S1_15B to fully desulfate the non-reducing terminal galactose. Tetrasaccharides with full occupation of the three residues at the non-reducing end could be modelled into electron density in the active sites of both monomers in the asymmetric unit (Fig. S3I). The recognition of the sugars involved a complex network of hydrogen bonds, including specific interactions with 2S modifications on the monosaccharides occupying the +1 and +2 subsites (Fig. 8B). Though the bound oligosaccharide is in a very similar position to that observed in GH110A, the +1 and +2 subsites are poorly conserved between GH110A and GH110B.

**Figure 8.**
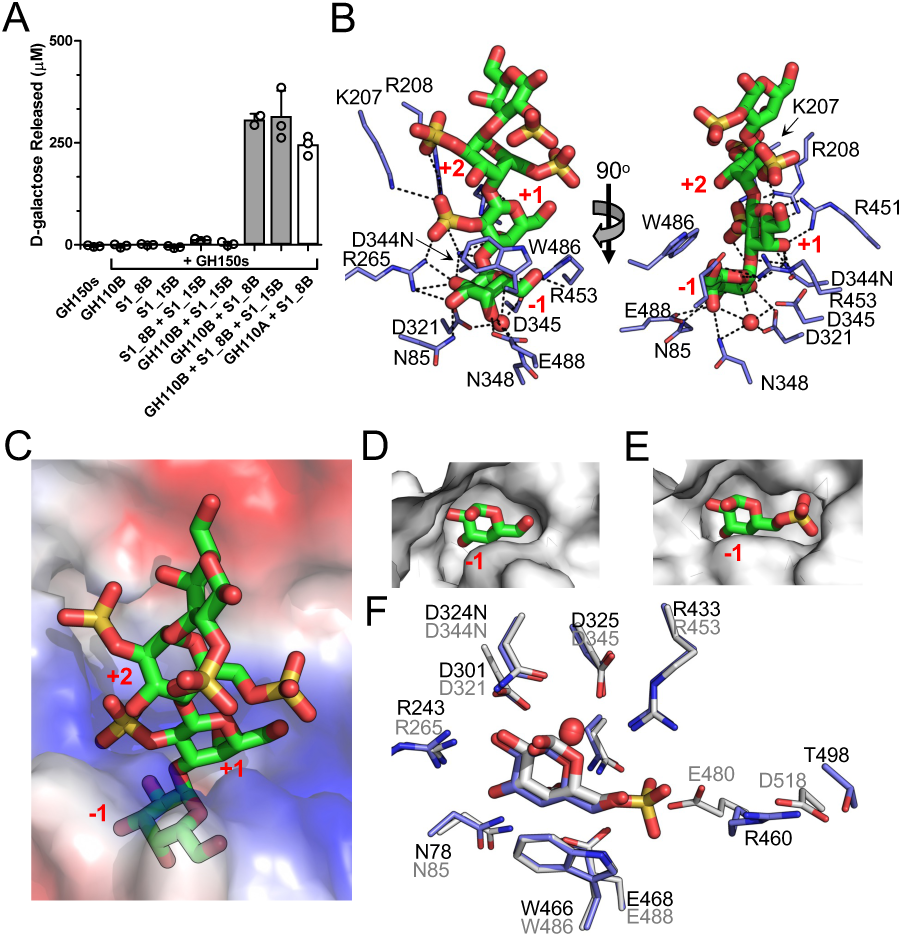
Characterisation of GH110B. A) Galactose release assay showing GH110B requires S1_8B and S1_15B pre-treatment of oligosaccharides before it could remove monosaccharides. Grey bars indicate the mean values of triplicate independent measurements and the error bars the standard deviation. Closed circles show individual data points. B) GH110B in complex with λ-NC4 that was pretreated with S1_8B and S1_15B to fully desulfate the non-reducing terminal galactose. The recognition of the sugars involved a complex network of hydrogen bonds, including specific interactions with 2S modifications on the monosaccharides occupying the +1 and +2 subsites. C) The unsulfated non-reducing terminal galactose is bound in -1 subsite at the bottom of a deep pocket. D) The surface of the -1 subsite closely complements the unsulfated galactose which is in contrast to E) the -1 subsite of GH110A which accommodates the 6S of G6S. F) Superimposition of GH110A and GH110B showing nearly identical -1 subsites apart from the region surrounding the 6S group of G6S.

There was no evidence in the electron density for sulfation of the non-reducing terminal galactose (Fig. S3I), which was bound in -1 subsite at the bottom of a deep pocket (Fig. 8C). As observed in our prior structures, the surface of the -1 subsite closely complements that of the unsulfated galactose (Fig. 8D), contrasting with the -1 subsite of GH110A, which clearly accommodates the 6S of G6S (Fig. 8E). The -1 subsites of GH110A and GH110B were nearly identical apart from the region surrounding the 6S group recognised by GH110A (Fig. 8F). In GH110B, the position occupied by R460 in GH110A is E480. R460 of GH110A is able to lean away from the active site, hydrogen bonding with the side chain of T498 and creating room for the 6S binding pocket. In contrast, the sidechain of E480 in GH110B is held in place by the sidechain of D518, blocking the 6S binding pocket. It is possible there are some allowable conformations of the E480/D518 pair in GH110B that allow accommodation of a terminal G6S residue, particularly if the 6S is oriented somewhat up and out of the pocket. This, combined with our quite long incubations in the activity assays, may explain the observed GH110B activity on a terminal G6S residue. We suggest, however, based on our structural observations that GH110B is optimally selective for a fully desulfated non-reducing terminal galactose residue. Therefore, GH110A is paired to work with S1_8B whereas GH110B is likely most efficient when functioning in concert with S1_8B and S1_15B.

### Identification of a monosaccharide specific galactose-2-sulfate-2-O-sulfohydrolase

By its membership in S1_8, we thought it likely that S1_8A would also be a 2-sulfate-2-O-sulfohydrolase. We determined its structure, which revealed a fold similar to S1_8B and conservation of the S and 0 subsites with S1_8B (Fig. 9A). The monosaccharide G2S was not available and we were unable to determine structures of S1_8A in complex with either galactose or galactosamine-2-sulfate (considered as a possible surrogate for G2S). To ultimately generate a complex, we hydrolyzed λ- NC4 with S1_8B, GH110A, S1_8C, and GH2 to release G6S (residue 1 and residue 3), galactose (residue 2), and G2S (residue 4 at the reducing terminus) (Fig. 10). We then soaked crystals of catalytically matured wild-type S1_8A with this mixture. The structure solved to 2.1 Å allowed modelling of sugars into the active sites of each of the two monomers in the asymmetric unit. In one active site, because the maturation state could not be clearly determined, the catalytic residue was modelled as the unmatured cysteine, while the bound monosaccharide was modelled as a single galactose residue (Fig. S3J) . The 2-OH of the galactose was oriented towards the catalytic machinery, consistent with 2S sulfatase specificity. In the other monomer, unbiased σ_a_-weighted F_o_-F_c_ omit maps generated by refinement prior to modelling the catalytic residue and bound ligand showed clear contiguous electron density joining the protein backbone and the galactose moiety (Fig. 9B). Having no other rational explanation for this unexpected finding, we modelled this complex as a covalent diester sulfo-galactose species, which has been proposed as an intermediate along one of the two hypothesized S1 sulfatase catalytic trajectories(28, 29). This intermediate could only be modelled based on G2S, again indicating 2S sulfatase specificity. Furthermore, in the active sites of both monomers, substrate binding results in the ordering of a loop that causes the active site to be enclosed to the extent that only a monosaccharide could be accommodated, revealing specificity on a G2S monosaccharide (Fig. 9C and 9D). We interpreted these structural observations as being most consistent with S1_8A being a monosaccharide specific galactose-2-sulfate-2-O-sulfohydrolase.

**Figure 9.**
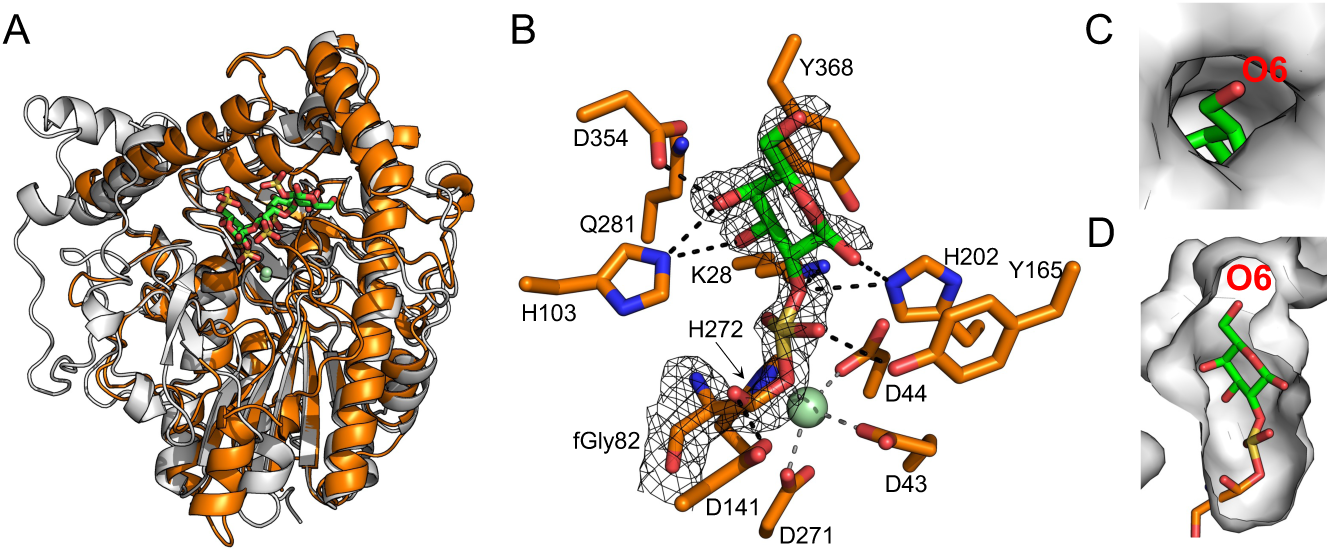
Characterisation of monosaccharide specific 2S sulfatase S1_8A. A) Superimposition of S1_8A and S1_8B showing overall structural fold conservation. B) Contiguous electron density between catalytically matured wild-type S1_8A catalytic residue fGly82 and bound G2S moiety modelled as covalent diester sulfo-galactose species, which is a proposed reaction intermediate. C and D) Binding of substrate results in movement of an active-site loop that causes the active site to be enclosed to the extent that only a monosaccharide could be accommodated during catalysis.

**Figure 10.**
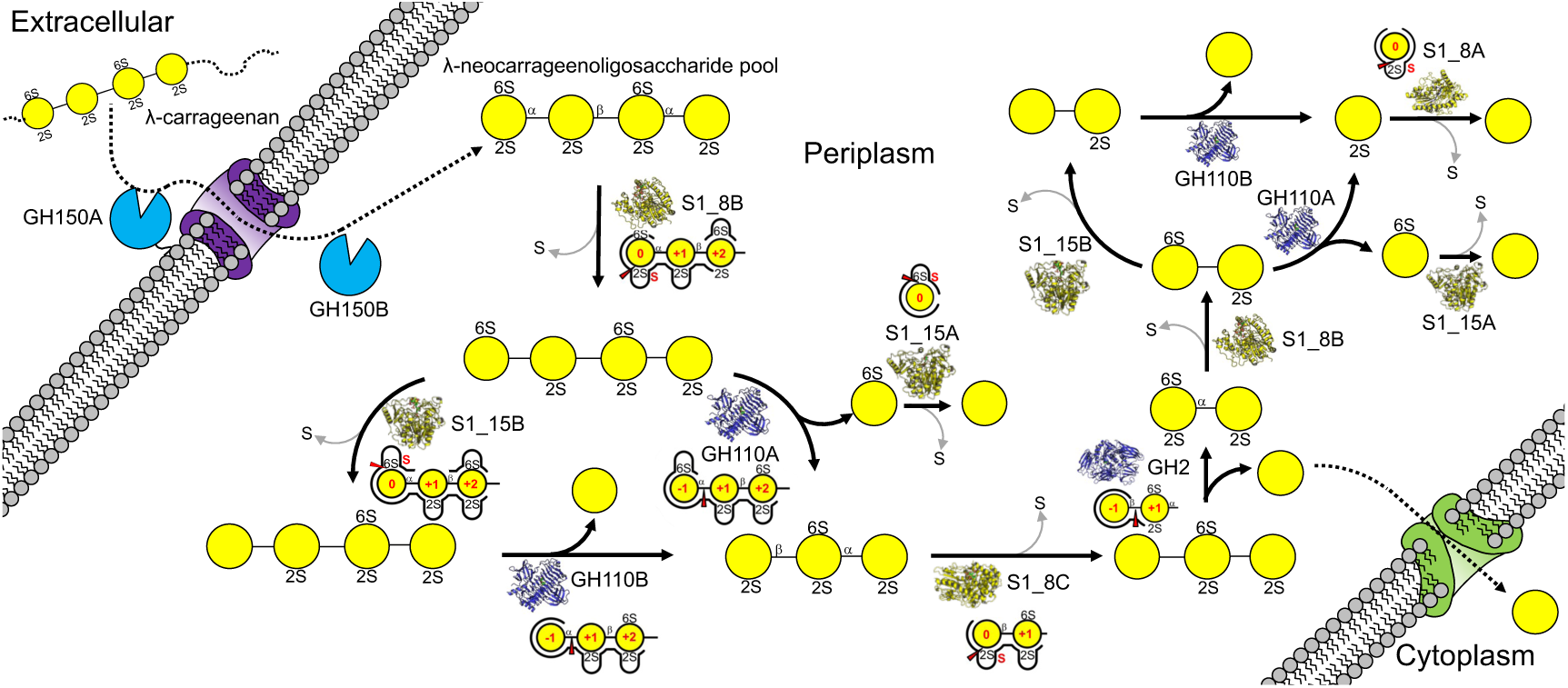
Schematic of λ-carrageenan metabolism by *Pseudoalteromonas distincta* U2A. Depolymerisation is initiated by the activity of the GH150 enzymes that produce a series of λ-neocarrageenoligosaccharides (joined yellow circles with sulfates indicated) with even numbered degrees of polymerization. λ-neocarrageenoligosaccharide breakdown in the periplasm proceeds via a cyclical process whereby the non-reducing end monosaccharides are sequentially removed by GH110A and GH110B and GH2. However, as each non-reducing terminal residue is revealed it must first be prepared for removal by specific sulfatase processing. GH110A requires removal of the 2-sulfate by S1_8 sulfatases. While this desulfation is sufficient to enable GH110B activity we suggest its optimal activity occurs after additional removal of the 6-sulfate by sulfatases from the S1_15 family. GH2 is active only when the terminal β-linked galactose is fully desulfated by the removal of the 2-sulfate group by S1_8C. The process of removing sulfates from the products of the cascade, namely the G6S released by GH110B and G2S left from the final hydrolysis step, to completely reduce λ-carrageenan to galactose (free yellow circle) is accomplished by sulfatases specific to G6S and G2S, *i.e.* S1_15A and S1_8A. Enzyme structures and schematics of the active sites representing the specificity of substrate recognition are provided for each reaction step.

## Discussion

The molecular details revealed by the biochemical and structural data presented in this work enable the proposal of a complex enzyme pathway that catalyzes the complete depolymerization of λ- carrageenan (Fig. 10). It is initiated by the activity of the GH150 enzymes that produce a series of λ-neocarrageenoligosaccharides with even numbered degrees of polymerization. λ- neocarrageenoligosaccharide breakdown in the periplasm proceeds via a cyclical process whereby the non-reducing end monosaccharides are sequentially removed by exo-α-1,3-galactosidases (GH110A and GH110B) and a β-1,4-galactosidase (GH2). However, as each non-reducing terminal residue is revealed it must first be prepared for removal by specific sulfatase processing. GH110A requires removal of the 2-sulfate by S1_8 sulfatases. While this desulfation is sufficient to enable GH110B activity we suggest its optimal activity occurs after additional removal of the 6- sulfate by sulfatases from the S1_15 family. GH2, the only β-galactosidase encoded by the λ−CarPUL, is active only when the terminal β-linked galactose is fully desulfated by the removal of the 2-sulfate group by S1_8C. The process of removing sulfates from the products of the cascade, namely the G6S released by GH110B and G2S left from the final hydrolysis step, to completely reduce λ-carrageenan to galactose is accomplished by sulfatases specific to G6S and G2S, *i.e.* S1_15A and S1_8A. The cyclical action of this cascade on oligosaccharides of any length derived from λ-carrageenan would enable the complete depolymerization of any product resulting from the endo-hydrolytic action of the GH150 enzymes, theoretically including those with an incomplete sulfation pattern.

We have previously presented the κ/ι-carrageenan metabolism pathways in U2A (10). Although the substrates for these pathways and the λ-carrageenan pathway reported here belong to the same algae polysaccharide family, there is not a single enzyme shared between these catabolic cascades, highlighting the high degree of enzyme specificity required in these processes. This is also true when we compare the λ-carrageenan pathway to the ι-carrageenan pathway described in *Zobellia galactanivorans*, with the exception of the last step where free galactose is released by GH2, which is a shared final step(11). Otherwise, the κ/ι-carrageenan metabolic pathways require distinct endo-acting GH16 or GH82 carrageenases to initiate depolymerization. The GH16 family contains a wide diversity of glycoside hydrolases that are active on a broad range of β-1,4 or β1,3 glycosidic bonds in various marine polysaccharides(30). In comparison, the GH150 and GH82 families contain glycoside hydrolases that are exclusively active on λ-carrageenan, or ι- carrageenan, respectively (20, 21, 31–36). Interestingly, GH16, GH82 and GH150 family enzymes do not share any significant structural similarity or evolutionary trajectories, emphasising the distinct separation in enzymatic requirements for the processing of different carrageen classes. This distinction is further observed when comparing the subsequent steps in both catabolic cascades. For the κ/ι-carrageenan pathways, subsequent depolymerization requires cyclic processing by 4S (S1_19, S1_7) and 2S (S1_NC and S1_17) sulfatases before an *exo*-acting GH167 and an unknown β-neocarrabiosidase (or GH127 and a GH2, as the case of κ-carrageenan) can act to release desulfated galactose and 3,6-anhydro galactose into central carbon catabolism. In the λ- carrageenan pathway the *exo*-acting 2S S1_8 and 6S S1_15 families sulfatases are required to act before (and after, in the case of S1_8A and S1_15A) the *exo*-acting GH110 enzyme(s) and GH2 can release free galactose. This highlights the completely distinct enzymatic cascades and incredible substrate specificity of the enzymes deployed in each of these pathways.

Our findings highlight the significant metabolic demand placed on heterotrophic bacteria that utilize marine sulfated polysaccharides as a nutrient source. Sulfatases in particular represent the unique enzymatic logic adapted by these specialized bacteria, such as U2A, to access the chemical energy locked in these complex biopolymers. Although sulfatases have gained increasing interest from researchers in the recent years due to their potential application in many industrial applications (37–40), our knowledge of the molecular details that underpin the important functions of these enzymes is limited. Catalysis by sulfatases in the fGly-dependent family (i.e. the S1 family) is proposed to proceed via two possible mechanisms (28, 29). In the first mechanism, the transesterification-elimination mechanism, the fGly residue exists in the geminal diol form. One hydroxyl group on fGly acts as a nucleophile and performs an S_N_2 attack on the sulfur atom of the sulfate, thus releasing the conjugate alcohol and forming a covalent fGly-sulfate enzyme adduct. This reacts to eliminate the sulfate and a hydration reaction regenerates the hydrated geminal diol form of the fGly residue. Based on the interpretation of existing data, this is the favored mechanism (28, 41). In the second proposed mechanism, the addition-hydrolysis mechanism, nucleophilic attack of a sulfate oxygen on the fGly in the aldehyde form results in a covalent sulfate diester intermediate. Subsequent hydrolysis of this intermediate releases the alcohol conjugate leaving the fGly-sulfate enzyme adduct, which reacts to eliminate the sulfate and regenerate the fGly residue. The complex of S1_8A with covalently bound substrate unexpectedly supports this second mechanism as we appear to have fortuitously trapped the covalent sulfate diester intermediate. In accordance with the proposed role of conserved S1 sulfatase catalytic residues, H202 is positioned to act as a catalytic acid to donate a proton to the oxygen in the ester linkage joining the carbohydrate leaving group and the enzyme bound sulfate. The sulfur adopts an unusual distorted pentagonal conformation, with the apex presented to a water filled cavity that can access bulk solvent when the loop over the active site is in its open conformation. This suggests the potential for nucleophilic attack by an activated water, though specific residues involved in activating the water are not immediately evident from the structure. In combination with the proposed role of H202 this would complete hydrolysis of the S-O bond, releasing the carbohydrate and leaving the enzyme fGly-sulfate adduct. This suggests that sulfatases may achieve hydrolysis of the S-O bond through multiple mechanisms, likely depending on the nature of the leaving groups, though we acknowledge this requires further investigation.

Carrageenans have been used extensively in the food industry for decades as gelling, stabilising and clarifying agents (2, 3) and are becoming an important source of sustainable next-generation high value products relevant to the food and pharmaceutical industry (1, 3, 37, 42). Recent work has highlighted the potential role that carrageenan-active enzymes can play in polysaccharide remodelling for industrial applications (37, 42, 43). However, more detailed biochemical characterization and optimization of these enzymes is critical to ensure their biocatalytic power in these processes are utilized to their full potential. The detailed biochemical interrogation of the enzymes reported here, may help in future endeavours of applying enzymatic logic to the development of algal based high value products. In summary, we determined and biochemically investigated for the first time, the complete metabolic pathway for λ-carrageenan utilisation by the marine bacteria *Pseudoalteromonas distincta* (strain U2A). Our findings highlight the highly specialized and complex enzymatic requirements necessary for complete degradation and utilization of one of natures most sulfated biopolymers.

## Experimental Procedures

### Growth of Pseudoalteromonas isolates on carrageenan

Growth of *Pseudoalteromonas* isolates on λ-carrageenan was assessed at 25 °C in a SpectraMax M5 plate reader at 600 nm as previously described (44). Quadruplicate overnight cultures of each strain were grown in Zobell broth from individual colonies, washed with minimal marine medium (MMM) (45) and resuspended in MMM containing no carbon source. Dialyzed and lyophilized λ-carrageenan (0.5% w/v) was prepared in MMM and filter sterilized prior to use. Washed cells were used to inoculate 100 μL cultures 1/50 in 96-well microplates, and sealed plates were incubated for 60 hours with shaking and OD readings every 20 minutes. Control wells contained uninoculated carrageenan/oligosaccharide, carbohydrate-free MMM inoculated with bacteria, and MMM containing 0.4% (w/v) galactose inoculated with bacteria.

### Detection of carrageenases in Pseudoalteromonas distincta U2A cultures

U2A was grown in 3 mL cultures of MMM containing 0.2% (w/v) λ-carrageenan for 60 hours as described above. Cells were pelleted at 13,000 rpm for 3 minutes and the supernatants retained. The pellets were briefly washed in fresh MMM, re-centrifuged, then lysed in 150 μL BugBuster® Protein Extraction Reagent (Novagen) at room temperature for 20 minutes. To remove contaminating carbohydrate originating from the BugBuster®, lysed cell pellets were dialyzed against binding buffer (20 mM Tris pH 8.0, 500 mM NaCl) overnight at 4 °C in 100 μL dialysis buttons (Hampton Research) fitted with a 3 kDa MWCO membrane. Carrageenan digests contained 10 μL culture supernatant or lysed cell pellet in 20-50 μL reactions. Samples were analyzed by FACE or C-PAGE as described below.

### Cloning and mutagenesis

The gene fragments encoding GH150A, GH150B, GH110A, GH110B, GH2, S1_15A, S1_15B, S1_8A, S1_8B, and S1_8C, without predicted signal peptides, were amplified from *Pseudoalteromonas distincta* U2A genomic DNA (see Table S2 for primer sequences) and ligated into pET28a between the NheI and XhoI restriction sites, except for GH150A and GH150B which were ligated into custom pET28a maltose binding-protein (MBP) fusion construct using an approach described previously (10).

All DNA amplifications were done using CloneAmp Hifi PCR Premix. Mutations were created by site-directed mutagenesis (QuikChange Site-Directed Mutagenesis kit) (Table S2). All constructs were sequence confirmed by bi-directional sequencing.

### Protein expression and purification

The expression plasmid for the GH150A-MBP fusion protein was transformed into *E. coli* SHuffle^®^ T7, GH150B-MBP into *E. coli* BL21 (DE3) Star, and GH2 into and *E. coli* Tuner^™^ (DE3). Expression cultures comprised 2 L LB broth containing 50 μg mL^-1^ kanamycin sulfate, which were grown at 30 °C with agitation at 180 rpm until the cell density reached OD600 of ∼0.5. The temperature was then dropped to 16 °C and recombinant protein production was induced with a final concentration of 0.5 mM IPTG and the cultures were incubated for an additional 16 h.

All S1 proteins were produced in the presence or absence of a co-produced recombinant formylglyicine generating enzyme (FGE). In brief, expression plasmids were co-transformed into *E. coli* BL21 (DE3) Star with the plasmid pBAD/myc-his A Rv0712 (FGE) (Addgene plasmid #16132), which encodes a FGE to promote sulfatase maturation, and expressed as described previously (28). Transformed bacteria were grown in LB broth containing 50 μg mL^-1^ kanamycin sulfate, 100 μg mL^-1^ ampicillin, 20 μM CaCl_2_, and 4 μM CuCl_2_ as described previously (10).

For all protein productions, cultures were centrifuged at 6300 x g for 10 min. Harvested cells of expressed GH150A-MBP and GH150B-MBP fusion proteins were chemically lysed by resuspension in 35 % (w/v) sucrose, 1 % (w/v) deoxycholate, 1 % Triton X-100, 500 mM NaCl, 500 mM D-sorbitol, 1 mM dithiothreitol (DTT), 1 mM ethylenediaminetetraacetic acid (EDTA), 10 % glycerol, 20 mM Tris (pH 8.0), 10 mg lysozyme, and 0.2 μg mL^-1^ DNase. All other proteins were chemically lysed by resuspension in the same buffer minus the D-sorbitol, DTT, EDTA, and glycerol. Resulting lysates were clarified by centrifugation at 16,500 x g for 30 min.

All S1 proteins, GH2, GH110A, and GH110B proteins were purified by applying the lysate supernatant to a nickel affinity chromatography column and eluted with 20 mM Tris-HCl (pH 8.0), 500 mM NaCl, with a stepwise increase in imidazole concentration of 5, 10, 15, 20, 40, 50, 100, and 500 mM. The GH150A-MBP and GH150B-MBP fusion proteins lysate supernatant was applied to amylose resin and washed with 20 mM Tris (pH 7.5), 250 mM NaCl, 500 mM D-sorbitol, 10 % glycerol, 1 mM EDTA, and 1 mM DTT. Bound protein was eluted with wash buffer containing 10 mM maltose. All samples containing the protein of interest as judged by sodium dodecyl sulfate polyacrylamide gel electrophoresis were concentrated with an Amicon ultrafiltration cell (EMD Millipore) with a 10 kDa MWCO. Proteins of interest were further purified using a HiPrep 16/60 Sephacryl S-200 HR size exclusion chromatography column in 20 mM Tris (pH 8.0) and 500 mM NaCl. The GH150A-MBP and GH150B-MBP fusion proteins were concentrated with an Amicon ultrafiltration cell with a 50 kDa MWCO and further purified using a HiPrep 16/60 Sephacryl S-500 HR size exclusion chromatography column in 20 mM Tris (pH 7.5), 500 mM NaCl, 500 mM D- sorbitol, 1 mM EDTA, and 1 mM DTT. Pure sample, of all proteins of interest, as again assessed by SDS-PAGE were pooled and concentrated. All proteins used for crystallography were further processed by cleaving the N-terminal His-tag via overnight incubation with thrombin in thrombin cleavage buffer 20 mM Tris-HCl (pH 8.0), 500 mM NaCl, and 2.5 mM CaCl_2_ prior to size exclusion chromatography, which was performed as described above.

### Enzymes assays

Following incubation of carrageenan with cellular fractions fluorophore-assisted carbohydrate electrophoresis (FACE) was performed using methods as described (28). Reactions for analysis by carbohydrate polyacrylamide gel electrophoresis (C-PAGE) were performed with 1 µM of enzyme and 0.25% λ-carrageenan in 20 mM Tris (pH 8) overnight at 37 °C. C-PAGE gels were run as described previously (46). Reactions monitored by the release of D-galactose and/or G6S from λ-carrageenan were prepared by digestion with enzyme cocktails containing 1 μM of each enzyme (in 50 mM MES (pH 6.5) 0.5 M NaCl at 25 °C for 20 h. D-Galactose and/or G6S was quantified in a 96-well plate by enzymatic detection as described previously (9). All assays were performed in triplicate.

### Oligosaccharide production, purification, and characterization

For oligosaccharide production, λ- carrageenan was resuspended in 20 mM (pH 7.5), 500 mM NaCl, and 500 mM D-sorbitol to a final concentration of 2 %. Excess GH150B-MBP was added and incubated at 37 °C for 72-96 hours. Samples were frozen, lyophilized, resuspended in 2 mL 50 % ethanol and spun down at 12,000 x g for 5 minutes to remove any undigested λ-carrageenan. Resulting supernatant was then passed over a size exclusion chromatography column containing Bio-Gel P-2 Gel (BioRad) in 20 mM ammonium carbonate. Fractions were analyzed for the presence and size of oligosaccharides by thin layer chromatography (TLC) as described previously (28).

Samples of purified oligosaccharide were prepared for HPLC-MS by NaBH4 reduction followed by solid-phase extraction (SPE) exactly as previously described (9). Reduced, de-salted samples were analyzed by quadrupole-time-of-flight (QToF) MS in negative electrospray ionization mode exactly as previously described (9).

For capillary electrophoresis with laser-induced fluorescence (CE-LIF) analysis, samples were desalted by solid phase extraction using 250 mg ENVICarb SPE cartridges (9). The organic eluates (50% acetonitrile; 0.1% trifluoroacetic acid; 4 x 550 μL) were concentrated for several hours on a Savant SpeedVac concentrator (Thermo Scientific) before they were pooled in a 200 μl tube and freeze-dried. The dried material was derivatized at room temperature for 16 h with 8-aminopyrene- 1,3,6-trisulfonate (APTS) and NaBH_3_CN (Sigma) exactly as previously described (47). CE-LIF was performed using a PA800 ProteomeLab (Beckman Coulter) equipped with an argon ion laser module. Electrophoresis was performed in positive ion mode (outlet = cathode) at a constant 30 kV using a 20 μm (internal diameter) x 37 cm fused silica capillary (Polymicro Technologies) using a background electrolyte (BGE) consisting of 140 mM sodium borate (Sigma), pH 10. The capillary was conditioned by rinsing (20 psi) with 1 M NaOH (2 min), water (3 min) and BGE (3 min) before hydrodynamically (0.5 psi for 10 s) injecting sample. Peak areas were manually integrated using 32 Karat software (Beckman-Coulter) and post-integration processing was performed using Microsoft Excel.

### Crystallization, diffraction data collection and processing

All crystals were grown at 18 °C by hanging drop vapour-diffusion with 1:1 ratios of crystallization solution and protein. Crystallization conditions are given Table S3. 25% (v/v) ethylene glycol was used as the cryo-protectant for all crystals, with the exception of the GH110A crystals where the crystallization condition is already a cryoprotectant. Diffraction data were collected on an instrument comprising a Pilatus 200K 2D detector coupled to a MicroMax-007HF X-ray generator with a VariMaxTM-HF ArcSec Confocal Optical System and an Oxford Cryostream 800. Data were integrated, scaled and merged using HKL2000 (48).

### Structure solution and refinement

The structure of GH110A in complex with G6S was obtained by soaking crystals of this protein in excess G6S. Phases were determined by molecular replacement using the structure of GH110B (PDB ID 7JW4) as a search model. An initial model was constructed by autobuilding using ARP/wARP (49). To obtain a substrate complex, crystals of in inactive GH110A_D324N mutant were soaked in an excess of λ−NC4 that was treated prior with S1_8B. A substrate complex of GH110B_D344N was obtained by soaking crystals in an excess of λ−NC4 that was treated prior with S1_8B and S1_15B.

The structure of S1_8B was solved by molecular replacement using PHASER (50). The search model comprised an ensemble of the carrageenan specific S1_19A, S1_19B, and S1_NC sulfatases from *P. fuliginea* PS47 (PDB ID codes: 6BIA, 6PRM, 6PT4) (10). BUCCANEER (51) was used to automatically build a model, which was finished by manual building with COOT (52) and refinement with REFMAC (53). The final S1_8B model was used to solve the structures of S1_8A and S1_8B. To generate a complex of S1_8B, crystals of the S1_8B C84S mutant were soaked in crystallization solution supplemented with an excess of λ−NC4. To generate a complex of S1_8C, crystals of the S1_8C C93S mutant were soaked with an excess of λ−NC4 that was treated prior with both S1_8B and GH110B. To generate a complex of S1_8A, crystals of this protein produced under conditions that would limit its maturation (i.e. in the absence of co-expressed FGE) were soaked with an excess of λ−NC4 that was treated prior with S1_8B, GH110B, S1_8C, and GH2, which would result in production of the monosaccharides galactose, G6S, and G2S.

An initial molecular replacement solution for S1_15A was obtained using the same ensemble of sulfatase structures as for S1_8B using PHASER; however, the phases were not of sufficient quality to enable model building. Therefore, this solution was input for a cycle of SAD with partial model phasing in PHASER to take advantage of ordered iodide atoms from the crystallization solution.

Density improvement with PARROT (also leveraging non-crystallographic symmetry) followed by automated model building with BUCCANEER resulted in a model that was used for a second round of SAD with partial model phasing. This resulted in phases that enabled ARP/wARP to automatically build a model of ∼90% completeness. The partial model of S1_15A was used to determine the structure of an S1_15B C84S mutant in complex with G6S by molecular replacement. A substrate complex of S1_15B C84S was generated by soaking crystals in an excess of λ−NC4 that was treated prior with both S1_8B. To generate a complex of S1_15A, crystals of the S1_15A C94S mutant were soaked with an excess of G6S.

The structure of GH2 in complex with galactoisofagomine (GIF) was obtained by soaking crystals of this protein in excess GIF. Phases were determined by molecular replacement using the structure of a GH2A from *Bacteroides uniformis* (PDB ID 5T98) as a search model. An initial model was constructed by autobuilding using ARP/wARP.

All models were finished by manual building with COOT and refinement with REFMAC and/or phenix.refine. Waters were added in COOT with FINDWATERS and manually checked after refinement. In all datasets, refinement procedures were monitored by flagging 5% of all observations as “free” (54). Model validation was performed with MOLPROBITY (55). All data processing and model refinement statistics and PDB ID accession codes are shown in Table S3.

## Data availability

The Protein Data Bank (PDB) accession codes for the structure coordinates and structure factors are 9BEH, 9BEV, 9BEU, 9BEY, 9BB9, 9BES, 9BBD, 9BEF, 9BBA, 9BEP, 9BAS, 9BAU, and 9BAV.

## Supporting information

This article contains supporting information.

## Funding and additional information

This research was supported by a Natural Sciences and Engineering Research Council of Canada Discovery Grant (FRN 04355).

## Author Contributions

C.J.V., A.G.H and A.B.B. designed research; C.J.V., A.G.H, J.K.H., S.S-W., B.P., B.M., B.E.M., L.M., and N. performed research; C.J.V., A.G.H., W.F.Z, and A.B.B. analyzed data; and C.J.V. and A.B.B. wrote the paper.

