## Supplementary material for "The complete λ-carrageenan depolymerization cascade from a marine Pseudoalteromonad revealed by structural analysis of the enzymes"

* Alisdair B. Boraston.

**This PDF file includes:**

Figures S1 to S5
Tables S1 to S3

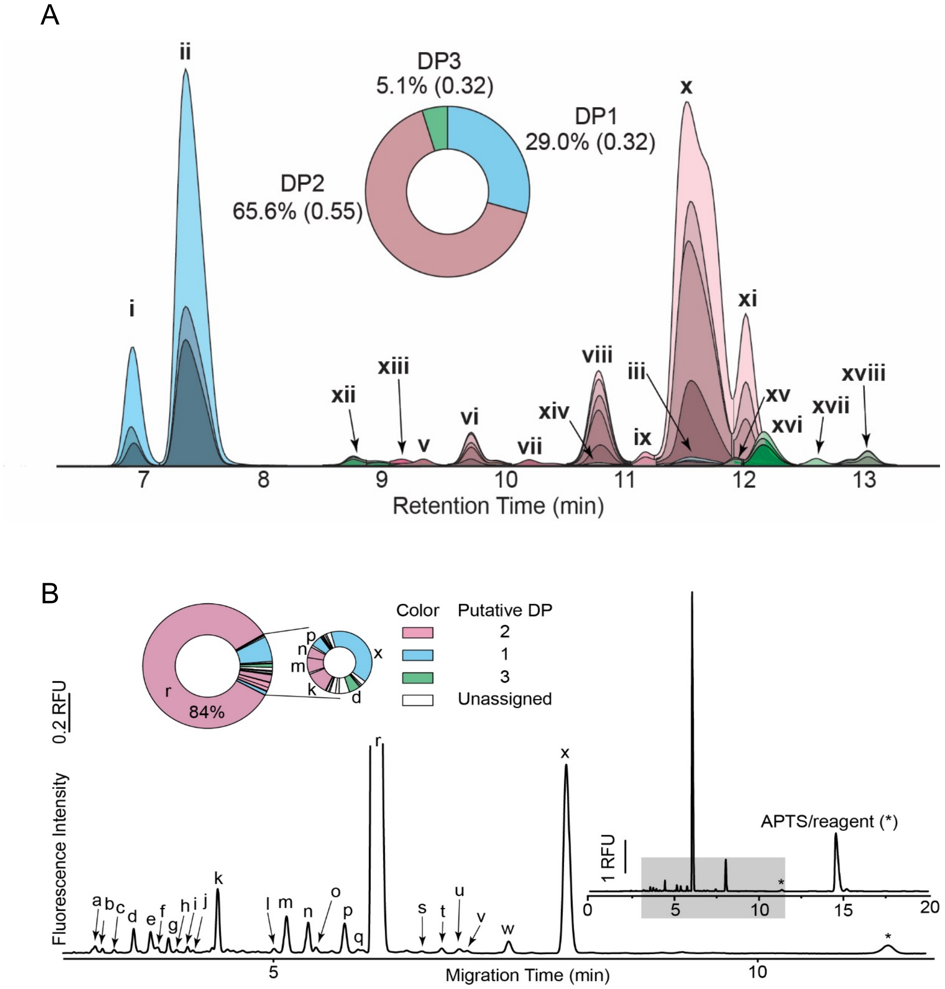

**Fig. S1. Analysis of λ-neocarrageenoligosaccharide.** A) HPLC-QToF-MS analysis of carrageenan oligomers. Oligosaccharide extracted ion chromatograms are color-coded based on their degree of polymerization (DP) with DP1 (blue) the basic lambda carrageenan repeating unit of two galactoses (hexoses; Hex) plus three sulfonate moieties. DP2 (purple) accounts for 66% of the total products detected, followed by DP1 (29%) and DP3 (green; 5%). The numbers in parentheses are the standard deviations of three technical replicates. A total of 18 unique oligosaccharides were detected. Oligosaccharide relative abundances were summed when peaks of identical DP and retention time exhibited obvious in-source de-*O*-sulfonation. Summary statistics are reported in Supplementary Table 1. B) Capillary electrophoresis (CE) analysis of digested λ-carrageenan reveals at least 24 products, ~84% of which is represented by a single oligosaccharide. It was hypothesized that the most abundant oligosaccharides detected by CE would directly correlate with the most abundant ions detected by HPLC-QToF-MS. Accordingly, a putative DP for the seven peaks of greatest area (r, p, n, m, k, d and x)—accounting for >96% of the total CE peak area—were assigned and color-coded as in panel A and Supplementary Table 1. In both panels the DP refers to the repeating disaccharide unit.

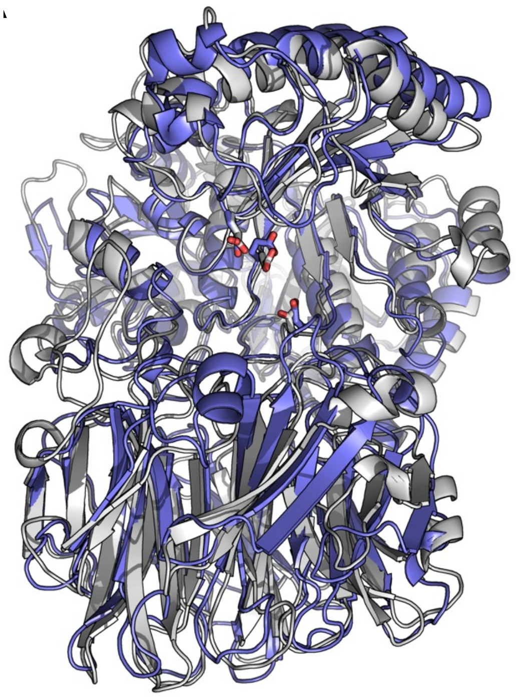

**Fig. S2. Comparison of GH150A (blue) and GH150B (grey) AlphaFold models.** Possible catalytic residues in the putative substrate binding cleft are shown as sticks.

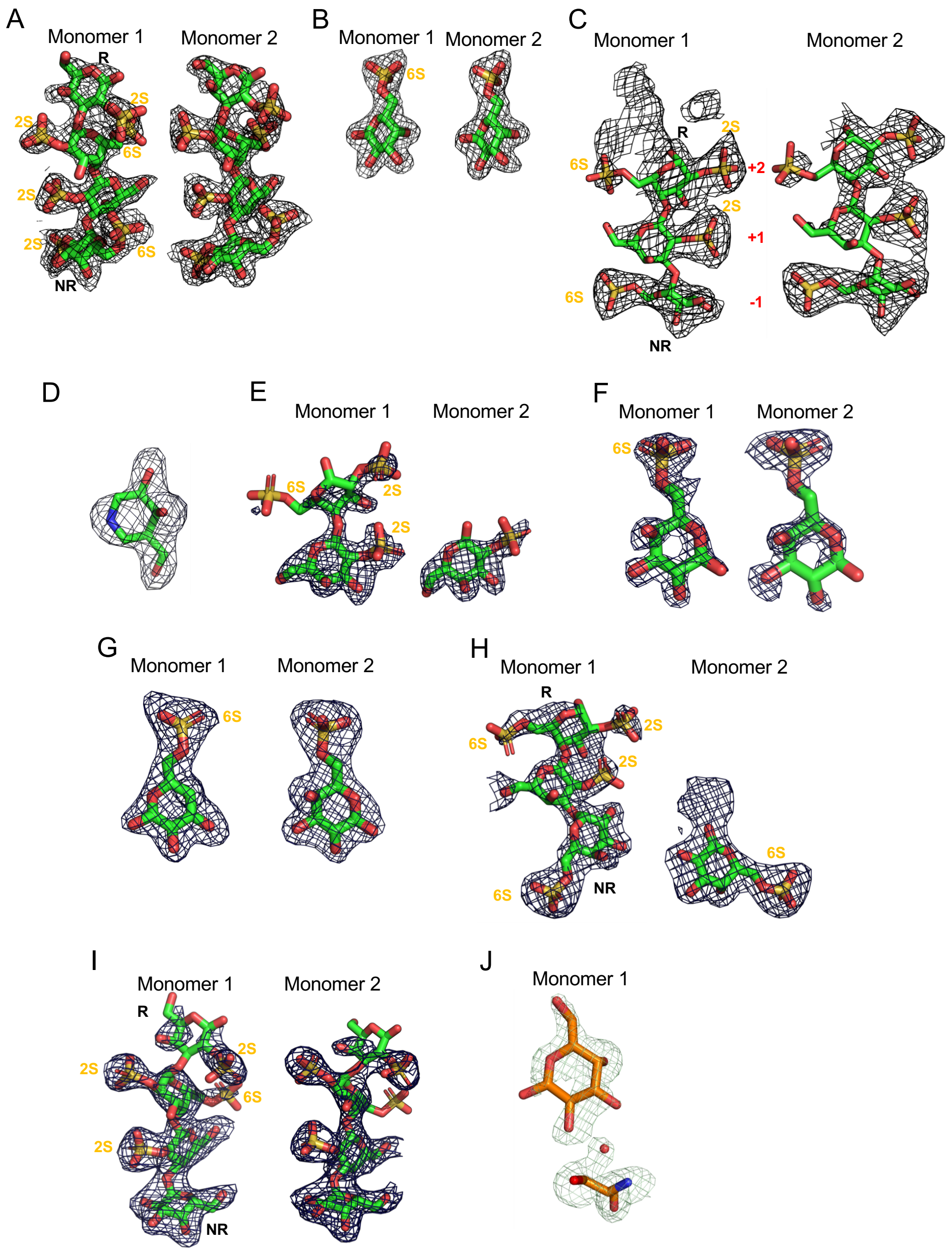

**Fig. S3. Electron density for bound substrates and ligands.** A) S1_8B C92S, B) GH110A with G6S, C) GH110A D324N with oligo, D) GH2 with galactoisofagomine, E) S1_8C C93S with oligo, F) S1_15A C94S with G6S, G) S1_15B C84S with G6S, H) S1_15B C84S with oligo, I) GH110B D344N with oligo, and S1_8A with oligo. The mesh represents $\sigma_a$-weighted $F_o$-$F_c$ maps generated prior to modelling the ligand. Maps are contoured at $3\sigma$.

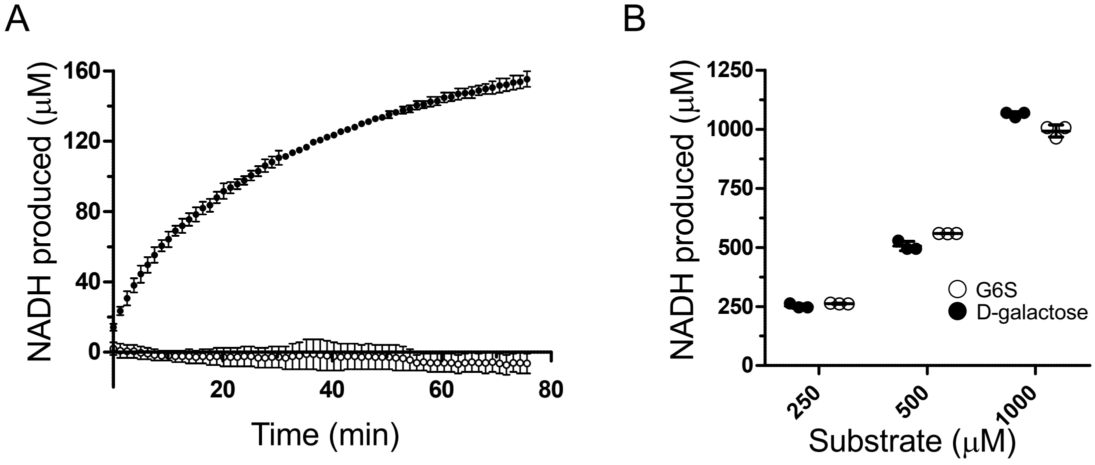

**Fig. S4.** Galactose release assay for A) GH110B (closed circles) and GH110A (open circles) on α-1,3-galactobiose. Data points show the mean and standard deviation of triplicate samples. B) Demonstration of equivalent detection of D-galactose and G6S. The mean and standard deviation of triplicate samples are shown with individual data points included.

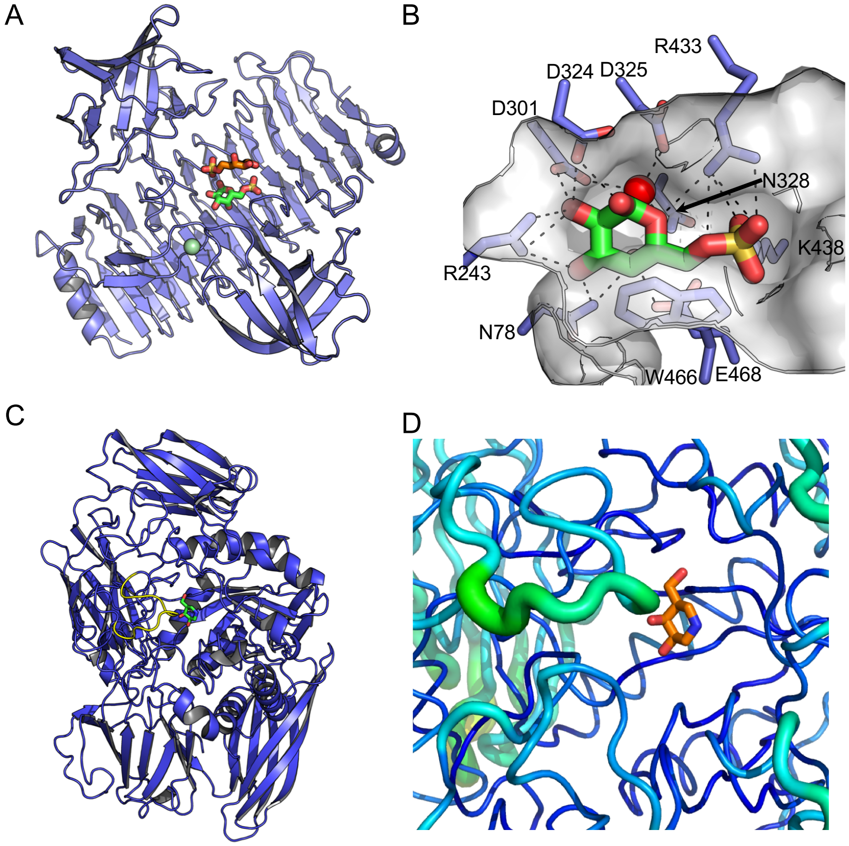

**Fig. S5. Structural analysis of GH110A and GH2.** A) Overall structure of GH110A bound to G6S. B) The active site of GH110A with G6S. C) Overall structure of GH2 bound to galactoisofagomine. The proposed flexible active site loop is coloured yellow. D) Tube representation of GH2 where the tube thickness and colour ramp from blue to green indicates increasing B-factor. The focused loop is that shown in panel C as yellow.

### Table S1. List of lambda carrageenan oligomers detected by full HPLC-QToF-MS.

| Peak | Composition | | | Formula | Retention Time[i] (Min) | | Area (%) | | Sum Area[ii] (%) | |
|---|---|---|---|---|---|---|---|---|---|---|
| | Hex | Sulfate[ii] (S) | Isobar | | Mean | SD | Mean | SD | Mean | SD |
| **Degree of polymerization = 1[iii]** | | | | | | | | | | |
| | 2 | 1 | | $C_{12}H_{24}O_{14}S_1$ | | | 0.7 | 0.10 | | |
| | 2 | 2 | | $C_{12}H_{24}O_{17}S_2$ | | | 2.7 | 0.04 | | |
| | 2 | 3 | | $C_{12}H_{24}O_{20}S_3$ | | | 0.4 | 0.02 | | |
| **i** | 2 | 3 | a | $C_{12}H_{24}O_{20}S_3$ | 6.90 | 0.020 | | | 4.0 | 3.89 |
| | 2 | 1 | | $C_{12}H_{24}O_{14}S_1$ | | | 4.2 | 0.21 | | |
| | 2 | 2 | | $C_{12}H_{24}O_{17}S_2$ | | | 5.8 | 0.03 | | |
| | 2 | 3 | | $C_{12}H_{24}O_{20}S_3$ | | | 13.9 | 0.13 | | |
| **ii** | 2 | 3 | b | $C_{12}H_{24}O_{20}S_3$ | 7.35 | 0.000 | | | 24.0 | 0.21 |
| | 2 | 1 | | $C_{12}H_{24}O_{14}S_1$ | | | 0.1 | 0.01 | | |
| | 2 | 2 | | $C_{12}H_{24}O_{17}S_2$ | | | 0.4 | 0.02 | | |
| | 2 | 3 | | $C_{12}H_{24}O_{20}S_3$ | | | 0.7 | 0.01 | | |
| **iii** | 2 | 3 | c | $C_{12}H_{24}O_{20}S_3$ | 11.55 | 0.038 | | | 1.2 | 0.04 |
| **iv** | 2 | 1 | a | $C_{12}H_{24}O_{14}S_1$ | 12.14 | 0.000 | 0.2 | 0.06 | | |
| **Degree of polymerization = 2** | | | | | | | | | | |
| **v** | 4 | 5 | a | $C_{24}H_{44}O_{36}S_5$ | 9.35 | | 0.2 | 0.07 | | |
| | 4 | 6 | | $C_{24}H_{44}O_{39}S_6$ | | | 0.2 | 0.02 | | |
| | 4 | 5 | | $C_{24}H_{44}O_{36}S_5$ | | | 0.5 | 0.07 | | |
| | 4 | 4 | | $C_{24}H_{44}O_{33}S_4$ | | | 0.7 | 0.04 | | |
| | 4 | 3 | | $C_{24}H_{44}O_{30}S_3$ | | | 0.5 | 0.01 | | |
| | 4 | 2 | | $C_{24}H_{44}O_{27}S_2$ | | | 0.7 | 0.02 | | |
| **vi** | 4 | 6 | a | $C_{24}H_{44}O_{39}S_6$ | 9.75 | 0.000 | | | 2.6 | 0.12 |
| **vii** | 4 | 5 | b | $C_{24}H_{44}O_{36}S_5$ | 10.23 | 0.000 | 0.1 | 0.08 | | |
| | 4 | 6 | | $C_{24}H_{44}O_{39}S_6$ | | | 0.6 | 0.02 | | |
| | 4 | 5 | | $C_{24}H_{44}O_{36}S_5$ | | | 1.9 | 0.06 | | |
| | 4 | 4 | | $C_{24}H_{44}O_{33}S_4$ | | | 2.4 | 0.02 | | |
| | 4 | 3 | | $C_{24}H_{44}O_{30}S_3$ | | | 1.6 | 0.02 | | |
| | 4 | 2 | | $C_{24}H_{44}O_{27}S_2$ | | | 2.6 | 0.02 | | |
| **viii** | 4 | 6 | b | $C_{24}H_{44}O_{39}S_6$ | 10.81 | 0.001 | | | 9.1 | 0.05 |
| | 4 | 3 | | $C_{24}H_{44}O_{30}S_3$ | | | 0.2 | 0.01 | | |
| | 4 | 2 | | $C_{24}H_{44}O_{27}S_2$ | | | 0.3 | 0.01 | | |
| **ix** | 4 | 3 | c | $C_{24}H_{44}O_{30}S_3$ | 11.20 | 0.011 | | | 0.5 | 0.01 |
| | 4 | 6 | | $C_{24}H_{44}O_{39}S_6$ | | | 0.3 | 0.05 | | |
| | 4 | 5 | | $C_{24}H_{44}O_{36}S_5$ | | | 3.9 | 0.14 | | |
| | 4 | 4 | | $C_{24}H_{44}O_{33}S_4$ | | | 11.6 | 0.20 | | |
| | 4 | 3 | | $C_{24}H_{44}O_{30}S_3$ | | | 10.8 | 0.19 | | |
| | 4 | 2 | | $C_{24}H_{44}O_{27}S_2$ | | | 19.5 | 0.11 | | |
| | 4 | 1 | | $C_{24}H_{44}O_{24}S_1$ | | | 0.2 | 0.01 | | |
| **x** | 4 | 6 | c | $C_{24}H_{44}O_{39}S_6$ | 11.56 | 0.017 | | | 46.1 | 0.38 |
| | 4 | 4 | | $C_{24}H_{44}O_{33}S_4$ | | | 1.1 | 0.04 | | |
| | 4 | 3 | | $C_{24}H_{44}O_{30}S_3$ | | | 2.2 | 0.08 | | |
| | 4 | 2 | | $C_{24}H_{44}O_{27}S_2$ | | | 3.8 | 0.16 | | |
| | 4 | 1 | | $C_{24}H_{44}O_{24}S_1$ | | | 0.1 | 0.03 | | |
| **xi** | 4 | 4 | a | $C_{24}H_{44}O_{33}S_4$ | 12.04 | 0.000 | | | 7.1 | 0.27 |
| **Degree of polymerization = 3** | | | | | | | | | | |
| | 6 | 9 | | $C_{36}H_{64}O_{58}S_9$ | | | 0.0 | 0.00 | | |
| | 6 | 8 | | $C_{36}H_{64}O_{55}S_8$ | | | 0.2 | 0.00 | | |
| **xii** | 6 | 9 | a | $C_{36}H_{64}O_{58}S_9$ | 8.98 | 0.000 | | | 0.2 | 0.01 |
| **xiii** | 6 | 6 | a | $C_{36}H_{64}O_{49}S_6$ | 9.16 | 0.000 | 0.2 | 0.04 | | |
| **xiv** | 6 | 8 | a | $C_{36}H_{64}O_{55}S_8$ | 10.17 | 0.000 | 0.1 | 0.01 | | |
| | 6 | 8 | | $C_{36}H_{64}O_{55}S_8$ | | | 0.0 | 0.00 | | |
| | 6 | 6 | | $C_{36}H_{64}O_{49}S_6$ | | | 0.2 | 0.01 | | |
| | 6 | 5 | | $C_{36}H_{64}O_{46}S_5$ | | | 0.1 | 0.00 | | |

| | | | | | | | | | | |
|---|---|---|---|---|---|---|---|---|---|---|
| | 6 | 4 | | $C_{36}H_{64}O_{43}S_4$ | | | 0.1 | 0.01 | | |
| **xv** | 6 | 8 | a | $C_{36}H_{64}O_{55}S_8$ | 11.9 | 0.008 | | | 0.3 | 0.09 |
| | 6 | 6 | | $C_{36}H_{64}O_{49}S_6$ | | | 0.9 | 0.03 | | |
| | 6 | 5 | | $C_{36}H_{64}O_{46}S_5$ | | | 1.0 | 0.04 | | |
| | 6 | 4 | | $C_{36}H_{64}O_{43}S_4$ | | | 0.7 | 0.04 | | |
| **xvi** | 6 | 6 | a | $C_{36}H_{64}O_{49}S_6$ | 12.12 | 0.000 | | | 2.3 | 0.58 |
| | 6 | 8 | | $C_{36}H_{64}O_{55}S_8$ | | | 0.2 | 0.00 | | |
| | 6 | 5 | | $C_{36}H_{64}O_{46}S_5$ | | | 0.1 | 0.02 | | |
| | 6 | 4 | | $C_{36}H_{64}O_{43}S_4$ | | | 0.0 | 0.00 | | |
| **xvii** | 6 | 8 | b | $C_{36}H_{64}O_{55}S_8$ | 12.73 | 0.167 | | | 0.3 | 0.03 |
| | 6 | 6 | | $C_{36}H_{64}O_{49}S_6$ | | | 0.4 | 0.20 | | |
| | 6 | 5 | | $C_{36}H_{64}O_{46}S_5$ | | | 0.4 | 0.01 | | |
| | 6 | 4 | | $C_{36}H_{64}O_{43}S_4$ | | | 0.2 | 0.04 | | |
| **xviii** | 6 | 6 | b | $C_{36}H_{64}O_{55}S_8$ | 13.06 | 0.001 | | | 1.1 | 0.04 |

**Notes: (i)** All data are reported as the mean (SD = standard deviation) of $N$ = 3 technical replicates. **(ii)** Frequent in-source de-*O*-sulfonation was observed as evidenced by oligosaccharides of identical degree of polymerization (DP) and retention time but with one or more sulfonate ($SO_3$) groups missing. The relative abundances of these de-sulfonated oligosaccharides are summed and recorded in the highlighted rows immediately following the list of de-sulfonated ions with the parent ion assumed to be the oligomer with the greatest number of sulfonates. Oligosaccharides where only a single ion was detected are considered unique and likewise highlighted. **(iii)** degree of polymerization refers to the number of repeated disaccharide units.

**Table S2:** Oligonucleotide primer sequences used for gene amplification

| | |
|---|---|
| MBP-GH150A FWD | CTGTACTTCCAGAGCTGCGTTATCCCTACTGTTG |
| MBP-GH150A REV | GTGGTGGTGGTGGTGCAATGTTGAACTTTGCATGTTTC |
| MBP-GH150B FWD | CTGTACTTCCAGAGCAGTGAAGTTTCACAAGATTATTTTACG |
| MBP-GH150B REV | GTGGTGGTGGTGGTGGTAACCAACATCAAAAAATCGGAC |
| PET28 MBP FWD | GCTCTGGAAGTACAGGTTCTC |
| PET28 MBP REV | CACCACCACCACCACC |
| GH110A FWD | CTAGCTAGCAAAGAGGTTTTAACTTTTG |
| GH110A REV | CCGCTCGAGTTACTTAATAGAGCCGTCGTC |
| GH110A D324N FWD | GAAAATATGCTAAATGACGGCGCAAACGTA |
| GH110A D324N REV | TACGTTTGCGCCGTCATTTAGCATATTTTC |
| GH2 FWD | CAGCCATATGGCTAGCAATGATGATAGAGTAAGCTTTAATAGTGGATGGTTATTC |
| GH2 REV | GGTGGTGGTGCTCGAGTTACTGGCTAACTAATACAACTGAACTTGAAATTAATTTTTC |
| S1_15A FWD | CGCGGCAGCCATATGGGGAAACTTACCAGTGATGATAAGAAAC |
| S1_15A REV | CCG CTCGAG TTACTTACGGGCTTTAA |
| S1_15A C94S FWD | CTGCTGCGACATCCACACCTTCTCGATATTCATTG |
| S1_15A C94S REV | GAGAAGGTGTGGATGTCGCAGCAGAACTGTG |
| S1_15B FWD | CGCGGCAGCCATATGGAGGCGCAAAATAGTGCG |
| S1_15B REV | TGGTGGTGCTCGAGTTCATTTTAGGCTCTTTAAACGTATAG |
| S1_15B C84S FWD | GCACACTCCTCTCCATCAAGATATTC |
| S1_15B C84S REV | GATGGAGAGGATGTTGCTGCA |
| S1_8A FWD | CTAGCTAGCCAAAGTGCTAGTGATAGT |
| S1_8A REV | CCGCTCGAGCTACTTGCCTTGC |
| S1_8B FWD | CTAGCTAGCAATGTAGAGGTGGACACC |
| S1_8B REV | CCGCTCGAGCTACTGAAATTTCTTTCGGTA |
| S1_8B C92S FWD | AACGTGTTGCAGAACTAACTGGAGCGGGCT |
| S1_8B C92S REV | CAGCCCGCTCCAGTTAGTTCTGCAACACGTT |
| S1_8C FWD | CTAGCTAGCGAGTCTTATGCTATATCG |
| S1_8C REV | CCGCTCGAGTTATATATTTTTCAAATAGT |
| S1_8C C93S FWD | GCCTGTATCCTCAACAGCAAGAAC |
| S1_8C C93S REV | CTGTTGAGGATACAGAATTAG |

**Table S3:** X-ray data collection and structure statistics

| | GH110A | GH110A_D324N | GH110B_D344N |
|---|---|---|---|
| | G6S | λ-oligo | λ-oligo |
| ***Data Collection*** | | | |
| Beamline | In-house | In-house | In-house |
| Wavelength | 1.541 | 1.541 | 1.541 |
| Space Group | C2 | C2 | C2 |
| Cell Dimensions | | | |
| *a, b, c* (Å) | 230.11, 77.48, 116.03 ($\beta$=113.39) | 231.51, 77.48, 116.04 ($\beta$=113.21) | 168.46, 128.35, 98.91 ($\beta$=122.15) |
| Resolution (Å) | 20.00-2.25 (2.29-2.25) | 30.00-2.60 (2.64-2.60) | 30.00-2.40 (2.44-2.40) |
| $R_{merge}$ | 0.110 (0.566) | 0.194 (0.535) | 0.117 (0.479) |
| $R_{pim}$ | 0.065 (0.373) | 0.089 (0.351) | 0.059 (0.332) |
| CC1/2 | 0.992 (0.815) | 0.975 (0.854) | 0.993 (0.803) |
| $\langle I/\sigma I \rangle$ | 10.6 (2.0) | 7.3 (1.7) | 11.5 (1.9) |
| Completeness (%) | 99.8 (99.8) | 99.8 (99.8) | 99.8 (99.9) |
| Redundancy | 3.6 (3.0) | 5.2 (3.3) | 4.7 (3.0) |
| No. of Reflections | 298,900 | 294,134 | 322,076 |
| No. Unique | 87,387 | 58,719 | 69,196 |
| ***Refinement*** | | | |
| Resolution (Å) | 2.25 | 2.60 | 2.40 |
| $R_{work}/R_{free}$ | 0.21/0.25 | 0.23/0.27 | 0.20/0.24 |
| No. of Atoms | | | |
| Protein | 4563 (A), 4571 (B) | 4549 (A), 4565 (B) | 4577 (A), 4569 (B) |
| Ligand | 30 (αG6S), 64 (βG6S) 2 ($Ca^{2+}$), 9 ($Cl^-$) | 100 (λ-NC3), 2 ($Ca^{2+}$) 7 ($Cl^-$) | 122 (λ-NC4), 43 ($I^-$) 24 ($Cl^-$), 28 (EDO) 13 (PG4) |
| Water | 488 | 368 | 212 |
| *B*-factors | | | |
| Protein | 37.8 (A), 35.3 (B) | 48.9 (A), 46.4 (B) | 40.9 (A), 40.5 (B) |
| Ligand | 30.2 (αG6S) 55.3 (βG6S) 33.0 ($Ca^{2+}$), 51.1 ($Cl^-$) | 68.0 (λ-NC3) 38.2 ($Ca^{2+}$), 56.6 ($Cl^-$) | 51.4 (λ-NC4), 70.1 ($I^-$) 44.5 ($Cl^-$), 42.6 (EDO) 45.6 (PG4) |
| Water | 37.9 | 46.8 | 38.6 |
| r.m.s.d. | | | |
| Bond Lengths (Å) | 0.005 | 0.003 | 0.008 |
| Bond Angles (°) | 0.794 | 0.632 | 0.991 |
| Ramachandran (%) | | | |
| Preferred | 95.6 | 94.5 | 94.9 |
| Allowed | 3.9 | 5.0 | 5.1 |
| Disallowed | 0.5 | 0.5 | 0 |
| PDB ID | 9BEH | 9BEV | 9BEU |

| | GH2 | S1_8A | S1_8A |
|---|---|---|---|
| | Galactoisofagomine | Native/MR | Complex |
| ***Data Collection*** | | | |
| Beamline | In-house | In-house | In-house |
| Wavelength | 1.541 | 1.541 | 1.541 |
| Space Group | $P2_1$ | P1 | P1 |
| Cell Dimensions | | | |
| *a, b, c* (Å) | 86.5, 70.3, 89.2 (β=113.5) | 50.03, 56.34, 97.20 (α=77.43, β=76.36, γ=63.91) | 49.68, 56.20, 97.23 (α=77.44, β=76.06, γ=64.03) |
| Resolution (Å) | 30.00-2.40 (2.44-2.40) | 30.00-1.85 (1.88-1.85) | 30.00-2.10 (2.14-2.10) |
| $R_{merge}$ | 0.121 (0.318) | 0.104 (0.396) | 0.101 (0.297) |
| $R_{pim}$ | 0.058 (0.231) | 0.046 (0.260) | 0.069 (0.146) |
| CC1/2 | 0.987 (0.821) | 0.995 (0.787) | 0.981 (0.915) |
| $<I/\sigma I>$ | 11.3 (2.5) | 14.2 (3.0) | 9.9 (2.3) |
| Completeness (%) | 97.9 (78.4) | 99.5 (97.9) | 95.0 (88.7) |
| Redundancy | 3.7 (2.0) | 5.0 (2.8) | 2.8 (2.2) |
| No. of Reflections | 142,147 | 398,197 | 142,346 |
| No. Unique | 38,258 | 78,931 | 51,285 |
| ***Refinement*** | | | |
| Resolution (Å) | 2.40 | 1.85 | 2.10 |
| $R_{work}/R_{free}$ | 0.19/0.23 | 0.16/0.19 | 0.20/0.26 |
| No. of Atoms | | | |
| Protein | 6415 | 3894 (A), 3857 (B) | 3869 (A), 3850 (B) |
| Ligand | 10 (GIF), 48 (EDO) | 2 ($Ca^{2+}$), 8 (EDO) | 12 (GAL), 2 ($Ca^{2+}$) 4 (EDO) |
| Water | 393 | 629 | 242 |
| *B*-factors | | | |
| Protein | 36.3 | 19.0 (A), 19.4 (B) | 24.1 (A), 24.5 (B) |
| Ligand | 32.0 (GIF), 39.4 (EDO) | 14.2 ($Ca^{2+}$) 33.6 (EDO) | 29.4 (GAL) 25.2 ($Ca^{2+}$) 25.6 (EDO) |
| Water | 36.5 | 23.4 | 22.5 |
| r.m.s.d. | | | |
| Bond Lengths (Å) | 0.002 | 0.010 | 0.007 |
| Bond Angles (°) | 0.540 | 1.077 | 1.169 |
| Ramachandran (%) | | | |
| Preferred | 96.3 | 96.6 | 95.0 |
| Allowed | 3.6 | 3.4 | 4.7 |
| Disallowed | 0.1 (1 residue S511) | 0 | 0.3 |
| PDB ID | 9BEY | 9BB9 | 9BES |

| | S1_8B | S1_8B C92S | S1_8C | S1_8C C93S |
|---|---|---|---|---|
| | Native/MR | λ-oligo | Native/MR | λ-oligo |
| ***Data Collection*** | | | | |
| Beamline | In-house | In-house | In-house | In-house |
| Wavelength | 1.541 | 1.541 | 1.541 | 1.541 |
| Space Group | $P2_12_12_1$ | $P2_12_12_1$ | $P2_1$ | I222 |
| Cell Dimensions | | | | |
| *a, b, c* (Å) | 93.84, 103.09, 148.16 | 80.31, 102.74, 190.07 | 156.90, 54.56, 159.29 ($\beta$=113.77) | 98.42, 162.13, 231.03 |
| Resolution (Å) | 30.00-2.30 (2.34-2.30) | 30.00-2.10 (2.14-2.10) | 30.00-2.20 (2.24-2.20) | 30.00-1.90 (1.93-1.90) |
| $R_{merge}$ | 0.103 (0.436) | 0.110 (0.237) | 0.088 (0.292) | 0.064 (0.474) |
| $R_{pim}$ | 0.053 (0.306) | 0.041 (0.213) | 0.043 (0.231) | 0.031 (0.362) |
| CC1/2 | 0.996 (0.869) | 0.996 (0.835) | 0.995 (0.824) | 0.997 (0.739) |
| $<I/\sigma I>$ | 18.5 (2.0) | 19.0 (3.2) | 13.5 (2.6) | 21.6 (1.6) |
| Completeness (%) | 96.1 (71.5) | 99.2 (92.1) | 98.0 (92.8) | 99.7 (99.0) |
| Redundancy | 6.8 (2.5) | 5.8 (2.1) | 4.2 (2.1) | 4.2 (2.4) |
| No. of Reflections | 425,774 | 528,434 | 513,831 | 608,054 |
| No. Unique | 62,174 | 91,025 | 123,433 | 144,283 |
| ***Refinement*** | | | | |
| Resolution (Å) | 2.3 | 2.10 | 2.20 | 1.90 |
| $R_{work}/R_{free}$ | 0.20/0.23 | 0.18/0.22 | 0.21/0.24 | 0.20/0.23 |
| No. of Atoms | | | | |
| Protein | 3653 (A), 3673 (B) | 4026 (A), 4037 (B) | 4866 (A), 4801 (B) 4749 (C), 4743 (D) | 4790 (A), 4792 (B) |
| Ligand | 40 ($SO_4^{2-}$), 16 (EDO) 2 ($Ca^{2+}$), 7 ($Cl^-$) | 138 (λ-NC4), 2 ($Ca^{2+}$) 2 ($Cl^-$), 15 ($SO_4^{2-}$) 8 (EDO) | 4 ($Ca^{2+}$), 4 (EDO) | 51 (λ-C2), 2 ($Ca^{2+}$) 4 ($Cl^-$) |
| Water | 356 | 656 | 421 | 698 |
| *B*-factors | | | | |
| Protein | 33.3 (A), 39.7 (B) | 27.6 (A), 26.0 (B) | 28.7 (A), 28.9 (B) 35.8 (C), 35.4 (D) | 20.2 (A), 25.6 (B) |
| Ligand | 56.8 ($SO_4^{2-}$) 44.1 (EDO) 22.7 ($Ca^{2+}$), 63.6 ($Cl^-$) | 55.7 (λ-NC4) 28.7 ($Ca^{2+}$) 47.2 ($Cl^-$), 54.2 ($SO_4^{2-}$) 43.2 (EDO) | 51.5 ($Ca^{2+}$) 34.4 (EDO) | 40.1 (λ-C2) 24.4 ($Ca^{2+}$), 17.3 ($Cl^-$) |
| Water | 37.0 | 30.4 | 28.9 | 26.0 |
| r.m.s.d. | | | | |
| Bond Lengths (Å) | 0.008 | 0.008 | 0.002 | 0.008 |
| Bond Angles (°) | 0.898 | 0.909 | 0.521 | 1.116 |
| Ramachandran (%) | | | | |
| Preferred | 97.0 | 97.4 | 97.6 | 98.0 |
| Allowed | 3.0 | 2.6 | 2.4 | 2.0 |
| Disallowed | 0 | 0 | 0 | 0 |
| PDB ID | 9BBD | 9BEF | 9BBA | 9BEP |

| | S1_15A | S1_15A C94S | S1_15B C84S | S1_15B C84S |
|---|---|---|---|---|
| | Native/I | G6S | G6S | λ-oligo |
| ***Data Collection*** | | | | |
| Beamline | In-house | In-house | In-house | In-house |
| Wavelength | 1.541 | 1.541 | 1.541 | 1.541 |
| Space Group | $P2_1$ | $P2_1$ | $P2_12_12_1$ | $P2_12_12_1$ |
| Cell Dimensions | | | | |
| *a, b, c* (Å) | 71.02, 106.79, 71.15 (β=105.38) | 71.87, 110.01, 71.79 (β=106.25) | 66.74, 93.85, 176.25 | 66.23, 92.87, 176.48 |
| Resolution (Å) | 30.00-2.25 (2.29-2.25) | 30.00-1.80 (1.83-1.80) | 30.00-2.30 (2.34-2.30) | 30.00-2.49 (2.54-2.49) |
| $R_{merge}$ | 0.063 (0.099) | 0.070 (0.368) | 0.091 (0.385) | 0.093 (0.326) |
| $R_{pim}$ | 0.024 (0.079) | 0.032 (0.198) | 0.047 (0.254) | 0.038 (0.154) |
| CC1/2 | 0.993 (0.962) | 0.996 (0.892) | 0.995 (0.785) | 0.997 (0.953) |
| $\langle I/\sigma I \rangle$ | 27.4 (7.3) | 19.1 (2.7) | 13.6 (2.4) | 19.1 (3.5) |
| Completeness (%) | 96.0 (69.0) | 99.8 (99.4) | 99.7 (98.9) | 98.8 (92.1) |
| Redundancy | 6.5 (1.8) | 3.8 (2.7) | 4.5 (2.7) | 6.2 (4.4) |
| No. of Reflections | 304,727 | 379,389 | 224,036 | 236,124 |
| No. Unique | 46,797 | 99,922 | 49,811 | 38,235 |
| ***Refinement*** | | | | |
| Resolution (Å) | | 1.80 | 2.30 | 2.49 |
| $R_{work}/R_{free}$ | | 0.19/0.24 | 0.19/0.24 | 0.22/0.26 |
| No. of Atoms | | | | |
| Protein | | 3719 (A), 3725 (B) | 3766 (A), 3767 (B) | 3784 (A), 3821 (B) |
| Ligand | | 32 (G6S), 2 ($Ca^{2+}$) | 32 (G6S), 2 ($Ca^{2+}$) 5 ($Cl^-$) | 50 (λ-NC3), 16 (G6S) 2 ($Ca^{2+}$), 3 ($Cl^-$) 8 (EDO) |
| Water | | 219 | 203 | 92 |
| *B*-factors | | | | |
| Protein | | 18.0 (A), 17.9 (B) | 31.8 (A), 36.0 (B) | 35.0 (A), 37.3 (B) |
| Ligand | | 15.7 (G6S), 27.2 ($Ca^{2+}$) | 28.0 (G6S), 32.1 ($Ca^{2+}$) 50.4 ($Cl^-$) | 57.8 (λ-NC3) 32.3 (G6S) 42.8 ($Ca^{2+}$), 54.9 ($Cl^-$) 45.4 (EDO) |
| Water | | 17.0 | 31.2 | 29.7 |
| r.m.s.d. | | | | |
| Bond Lengths (Å) | | 0.008 | 0.008 | 0.005 |
| Bond Angles (°) | | 1.177 | 0.960 | 0.807 |
| Ramachandran (%) | | | | |
| Preferred | | 94.9 | 97.0 | 96.3 |
| Allowed | | 5.1 | 3.0 | 3.7 |
| Disallowed | | 0 | 0 | 0 |
| PDB ID | | 9BAS | 9BAU | 9BAV |
